# Gut microbiota and maternal immune transfer at birth influence pre-allergic clinical outcome

**DOI:** 10.1101/2023.04.25.537372

**Authors:** Remy Villette, Djelika Traore, Elise Dhilly, Pierre Foucault, Eleonore Parisel, Delphine Sauce, Guy Gorochov, Gilles Kayem, Marta Schuhmacher, Isabella Annesi-Maesano, Martin Larsen, EarlyFOOD study group

**Affiliations:** Sorbonne Université, INSERM U1135, Centre d’Immunologie et des Maladies Infectieuses (CIMI-Paris), 75013, Paris, France; Département d’immunologie, Assistance Publique Hôpitaux de Paris (AP-HP), Hôpital Pitié-Salpêtrière, 75013 Paris, France; Trousseau Maternity Hospital, Assistance Publique Hôpitaux de Paris (AP-HP), Sorbonne Université, Paris, France; Environmental Engineering Laboratory, Departament d’Enginyeria Quimica, Universitat Rovira i Virgili, Tarragona, Spain; Institut Pierre Louis d’Epidémiologie et de Santé Publique, Equipe EPAR, Sorbonne Université, Paris, France; Institute Desbrest of Epidemiology and Public Health, INSERM and Montpellier University, Department of Allergic and Respiratory Diseases, Montpellier University Hospital, Montpellier, France; Institute of Clinical Physiology (IFC), National Research Council (CNR), Pisa, Italy; Department of Microbiology and Systems Biology, TNO Healthy Living and Work, Leiden, The Netherlands; Department of Preventive Dentistry, Academic Center for Dentistry Amsterdam, University of Amsterdam and Vrije Universiteit Amsterdam, Amsterdam, The Netherlands; Institute of Translational Pharmacology (IFT), National Research Council (CNR), Palermo, Italy

**Keywords:** Infant, microbiota, meconium, allergy, secretory IgA, IgAseq, 16S rRNA gene sequencing

## Abstract

The gut microbiota of 2-3 month-old infants is associated with later pre-allergic signs, while the microbiota at the time of allergic manifestation is not. We hypothesized that the infant gut microbiota and immune system are primed shortly after birth, and that this is influenced by maternal transfer of humoral immunity. We investigated the association between allergic outcomes and composition and humoral immunity to gut microbiota at birth, 2 months, and 2 years-of-age. Meconium microbiota clustered into three groups dominated by *Escherichia*, *Enterococcus*, and mixed genera, respectively. The *Escherichia* cluster was associated with protection against later allergic manifestations. We moreover studied the proportion and specificity of humoral immunity to gut microbiota. Humoral immunity to gut microbiota at birth was associated with future allergies. Future studies should evaluate whether interventions to alter gut microbiota and humoral immunity in early-life protects against allergy.

## Introduction

Microbiota is an important actor of human health, despite being well studied in adults, little is known about the early stages of microbial colonization after birth. Timing of the first colonization of neonates is controversial. Two competing theories describe initial gut microbiota colonization in humans; either colonization occurs *in utero*^1^ or alternatively following placental barrier breach during labor or C-section^1^. Recent work shows the importance of breastmilk, skin and feces bacteria transferred by the mothers to the infant when seeding the neonate ^3^. However, a large amount of the early colonizers are not found in any of the mother body sites indicating alternative sources of bacteria ^3, 4^. Nevertheless, birth route ^5–7^ and breastfeeding ^8, 9^ are implicated in early-life microbiota colonization and have been associated with later clinical outcomes^10^. Vaginal delivery and breastfeeding have been demonstrated to be protective against numerous diseases amongst which asthma^7^, allergic diseases^5, 6, 11^, obesity^12–14^, children infections^15, 16^, type-1 diabetes^17^ and auto-immune diseases^18^. These observations might be explained by a delayed establishment of key taxa in C-section born infants^19–21^ interfering with the normal maturation of the immune system ^22^ and a deregulated metabolism^23^. Vaginally born infants are colonized by *Escherichia*, *Staphylococcus, Bifidobacterium* and *Lactobacillus*^19, 24, 25^ while C-section born infants harbor more *Enterococcus, Streptococcus* and *Klebsiella* in their stools at birth^21^. Infant gut microbiota will transition from a low diversity microbiome in the first weeks, to a transitional *Bifidobacterium* dominated microbiota due to milk consumption, which shifts toward an adult-like microbiota when solid food is introduced. ^24^

Breast milk is a complex liquid composed of dietary fat, simple saccharides and complex saccharides, proteins, enzymes, immune cells, cytokines, growth factors, hormones and antimicrobial molecules^26^. A large proportion of the proteins transferred in the breast milk are immunoglobulins, allowing a vertical transfer of immunity from the mother toward the infant. Secretory IgA (sIgA) is the main antibody isotype transferred via breast milk with concentration at a peak in the colostrum (first breast milk) and declining over the first months postpartum ^27, 28^. Secretory IgA is actively secreted across various epithelial barriers via the poly Ig receptor (pIgR) mediated mechanism^29^ and contributes to the control of microbial colonization ^8, 30, 31^.

IgA+ B cells are detected in the neonate lamina propria 2 weeks postpartum, as well as secretory component, showing a sIgA production capacity.^32^ Despite this, infants have reduced sIgA secretion capacity and are dependent on the mothers input of sIgA at birth and during the first months of life (breastfeeding)^33^. Maternally transferred immunity confers protection against pathogens in early-life^8^. While the relation between sIgA secretion and microbiota is still poorly understood it’s been well shown that IgA can control pathogens and pathobionts via growth enchainment^30^, reduction of motility^34, 35^. Beyond the immune-exclusion mechanisms mentioned, sIgA also encompasses immune-inclusion mechanisms notably through the creation of ecological niches favoring the growth of slow-growing mutualists^31^ and shifting bacterial metabolism towards nutrient digestion^37^. In short, sIgA is capable of reducing pathogenicity and fitness of pathogens and pathobionts while increasing fitness of mutualists.

Early-life exposure to infectious events ^38, 39^ and allergens ^40, 41^ have been demonstrated to protect from later allergic manifestations. In addition, several studies have demonstrated an association between gut microbiota composition in the first months of life and allergic manifestations later in life^10, 42^. The effects were associated with a decrease in Veillonella or Lachnospira bacteria and an increase in diHOME metabolites. It therefore seems that the risk of developing an allergy is tightly linked with environmental exposure in early life.

Here we hypothesize that the first months of life are determinant for immune priming and memory development as well as later clinical outcome. Additionally, we hypothesize that sIgA is involved in the regulation of microbial colonization. For the first time we investigate the relationship between initial microbial colonization (meconium microbiota composition), host immunity and development of pre-allergic symptoms in the first year of life. We found that the composition and antibody-microbiota phenotype of gut microbiota at birth is associated with the development of infantile asthma, wheezing and allergies in total.

## Results

### Meconium microbiota has low alpha diversity and can be categorized into 3 distinct Meconium Community Types (MeCT)

Meconium samples have been analyzed by V3-V4 16S rRNA gene analysis to determine the microbiome composition. Meconium microbiota alpha diversity was generally much lower than adult microbiota alpha diversity but showed large disparity between samples (median=1.05 [0.05-3], maternal alpha diversity = 3.6, Supplemental figure 2A), additionally, the observed richness was between 1 and 66 ASVs detected (median=11.50 ASVs) compared to maternal gut microbiota (median=95 ASVs)(Supplemental figure 2B). Alpha diversity variation was not explained by sex, birth route or breastfeeding (Figure 1A) or any other clinical factor tested, such as maternal allergies, maternal vaginal infection or maternal BMI (Supplemental table 1). Meconium microbiota were dominated by a handful of taxa, *Escherichia-Shigella*, *Enterococcus*, *Staphylococcus*, *Bifidobacterium* and *Bacteroides*, which were detected in 87%, 81%, 64%, 54% and 51% of the samples, respectively. These bacteria represented more than 10% of the relative abundance in 59%, 39%, 15%, 18% and 13% of the samples, respectively (Figure 1B, Supplemental figure 2C). Meconium microbiomes were frequently composed almost entirely of one taxa. Eighty-nine samples (41%) were dominated with more than 80% relative abundance attributed to one taxa (ASV) (Figure 1B and Supplemental figure 2C). *Escherichia*-*Shigella* was higher in breastfed infants (Mann-Whitney Wilcoxon test, p=0.053) and vaginally delivered infants (Mann-Whitney Wilcoxon test, p=0.0016), while *Enterococcus* abundance was significantly higher in C-section delivered infants (Mann-Whitney Wilcoxon test, p=0.025). C-section born infants had a tendency to be more colonized by *Staphylococcus* (Figure 1C). Beta-diversity analysis was very consistent across different types of distance metrics (Supplemental figure 2D). Bray-Curtis dissimilarity was selected for the remainder of our analysis. Principal component analysis showed an effect of both breastfeeding and birth route on meconium microbiota composition (PERMANOVA, p=0.04 and 0.02 respectively, Figure 1D). These differences seemed to be driven by the *Escherichia/Shigella*, *Enterococcus* and *Staphylococcus* genus (Figure 1C). However, most of the beta-diversity variance was not explained by perinatal factors. We then performed a non-supervised hierarchical clustering and clustering validation (see Material and methods), which identified 3 distinct clusters based on meconium microbiome composition alone. We coined them Meconium Community Types (MeCT, Figure 1E). MeCT 1 was clearly dominated by *Escherichia-Shigella* genus, MeCT 2 was not dominated by any taxa and was very variable between samples, finally, MeCT 3 was clearly dominated by *Enterococcus* genus (Figure 1F).

**Figure 1.**
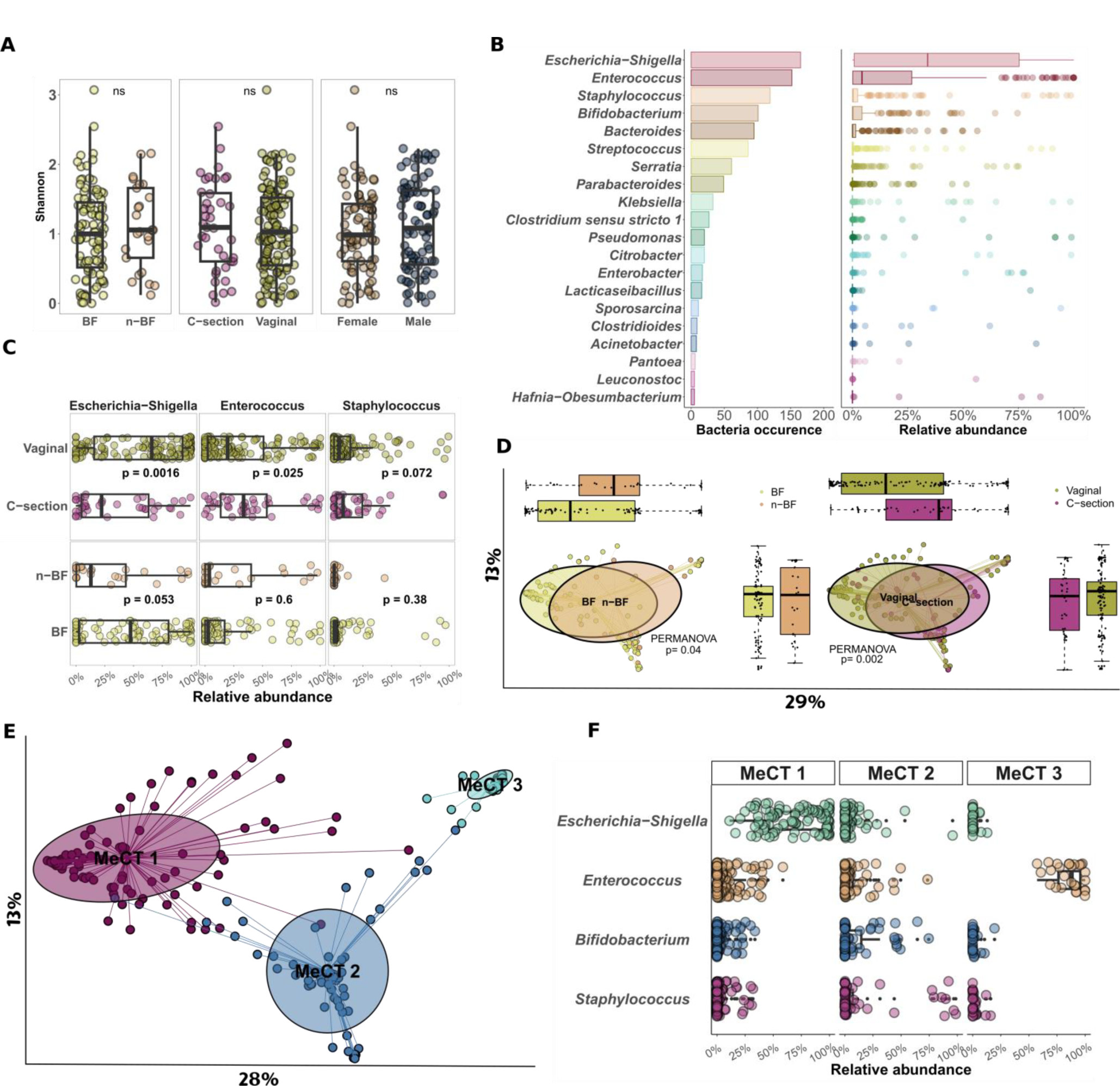
Meconium microbiota was sparse and segregated into three microbial community types. **A.** Shannon alpha diversity for breastfeeding, birth route and gender. **B.** Top 20 genera encountered in the meconium microbiota depicted as bacteria incidence (number of samples containing a given genus out of 187 samples) and the relative abundance for each sample. **C.** Differential abundance of the three most abundant taxa according to breastfeeding and birth route. **D.** Beta-diversity of meconium samples using a bray curtis dissimilarity matrix computed with a PCoA and assigned according to breastfeeding and birth route. Boxplots represent the sample distribution for each axis. **E.** PCoA based on Bray-Curtis dissimilarity and segregated by hierarchical clustering depicting three clusters designated Meconium Community types (MeCT). **F.** Relative abundance of the top 4 genera in meconium microbiota stratified by MeCT. All statistical tests conducted are Mann-Whitney Wilcoxon tests and PERMANOVA with 1000 permutations.

### Meconium community type and formula feeding are associated with clinical burden at 1 year of age

We next investigated if meconium microbiota composition was associated with pre-allergic clinical outcome at 1 year of life. Interestingly, we found an association between meconium composition and allergies at 1 year of life (PERMANOVA p=0.009, Figure 2A). To understand the microbial drivers of this result we then tested the association between meconium community types and clinical outcome at 1 year. The community types were associated with infantile asthma, infantile wheezing, lactose intolerance and overall allergies at 1 year (Mann-Whitney Wilcoxon test, p-values of 0.04, 0.03, 0.01 and 0.03, respectively, Figure 2B). MeCT 3 (*Enterococcus*) was associated with infant wheezing and infant asthma at 1 year (Figure 2C), while MeCT 2 (Mixed) was associated with lactose intolerance (Chi square test, p-value= 0.01, Supplemental figure 3A) and food allergy at 1 year (p-value= 0.06, Chi square test, Supplemental figure 3B). In fact lactose intolerance was specific to the MeCT 2 (Supplemental figure 3A). Interestingly, the MeCT 1 (*Escherichia*-*Shigella*) was associated with reduced incidence of pre-allergic clinical outcome at 1 year of life (p-value= 0.03, Figure 2D). We then looked at the link between the MeCT distribution and perinatal factors, such as birth route and feeding status. Meconium microbiota from children born vaginally were more frequently MeCT 1 while infants born by C-section were more frequently MeCT 2 and 3. Similarly, meconium microbiota from breastfed infants were more frequently MeCT 1 whereas formula fed infants were more frequently MeCT 2 and 3 (Figure 2D). *Escherichia* genus was increased in breastfed and vaginally delivered children and linked with reduced incidence of pre-allergic clinical outcome at 1 year. Finally, to assess the potential protective effect of breastfeeding we associated feeding status with clinical outcome at 1 year. Formula feeding was associated with a higher antibiotic usage and otitis during 1st year of life (Chi square test, p-value=0.009 and 0.017, Figure 2F and Supplemental figure 3 C-D). Of note, feeding status and birth route were not associated with later allergic manifestations but bronchitis was more present in infants with pre-allergic symptoms (Supplemental table 2).

**Figure 2.**
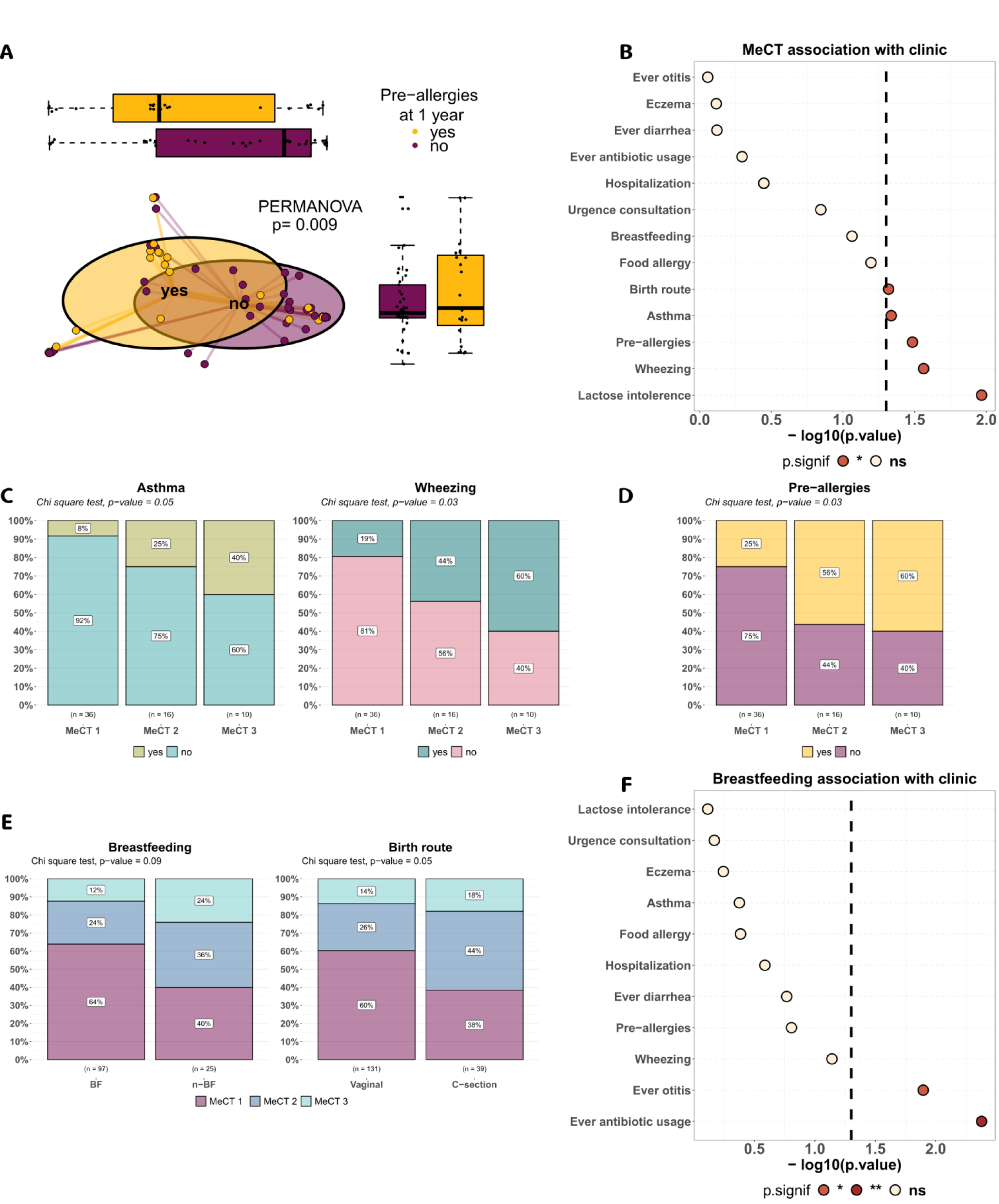
Microbiota composition at birth was associated with pre-allergic outcome at 1 year. **A.** Beta-diversity analysis of meconium microbiota segregated by allergies developed at 1 year of life. PCoA on Bray-Curtis dissimilarity, Axis 1: 32.1%, Axis 2: 14.5%, PERMANOVA test with a 1000 permutations. **B.** Association between perinatal factors and clinical outcomes at 1 year and MeCT. **C.** Association between infantile asthma and wheezing depending on the MeCT. **D.** Association between MeCTs and pre-allergy signs, including infantile asthma, wheezing and food allergy. **D.** Association between MeCT and breastfeeding (left panel) as well as birth route (right panel). **F.** Association between breastfeeding and clinical outcome at 1 year of age. All statistics are performed using a Chi Square test unless stated otherwise.

### Gut microbiota of infants at 2 years of age approach maternal gut microbiota composition

Of the 187 infants included at birth, 68 (36%) infants were lost to follow-up in part due to the COVID pandemic. 119 families responded to the 1 year follow up questionnaire. We retrieved 52 child stool samples and 41 matched maternal stool samples corresponding to 2 years follow up after birth. We subjected the gut microbiota composition to alpha and beta-diversity analysis comparing infant gut microbiota at different time points as well as maternal gut microbiota. Bray-Curtis dissimilarity analysis showed that the gut microbiome from 2 years-old children was clearly distant from the meconium microbiome and had approached the maternal gut microbiota (Figure 3A). Of note, despite the reduced distance to the maternal gut microbiota, the 2 year-old child gut microbiota remained significantly different from the maternal gut microbiota (PERMANOVA, p-value= 0.001, Figure 3B).

**Figure 3.**
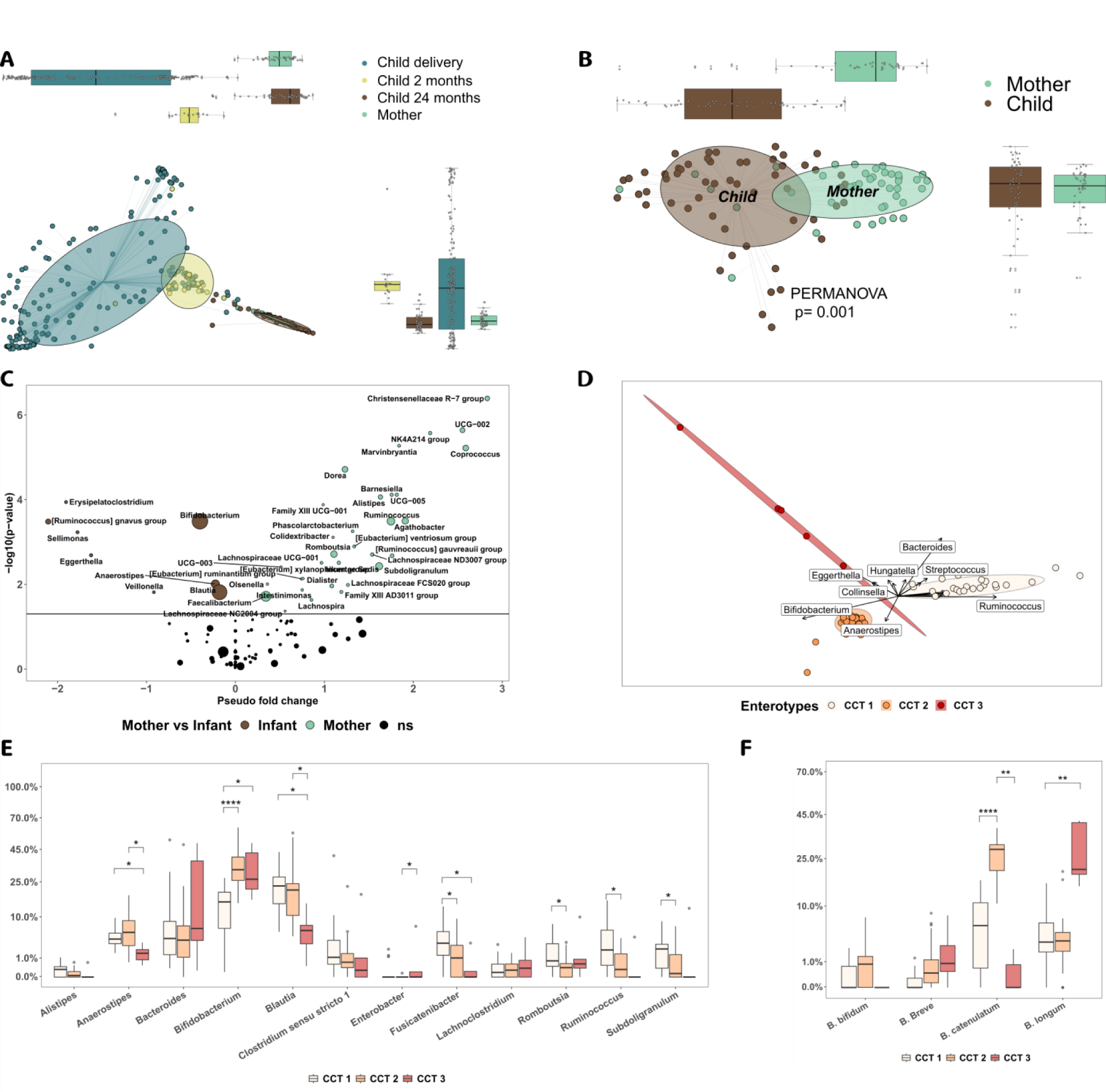
Child microbiomes gained maturity but remained less mature than the maternal microbiomes. **A.** Beta diversity analysis of gut microbiota from all points in time are depicted using a Bray-Curtis dissimilarity distance followed by a PCoA, boxplots represent the data distribution of each axis, here Axis 1: 20% and Axis 2: 12%. **B.** Beta-diversity analysis of mother versus child (24 months-of-age) based on Bray-Curtis dissimilarity and PCoA. **C.** Differential abundance analysis represented as a volcano plot. Taxa in green are enriched in maternal samples while brown taxa are enriched in child samples. **D.** Child Community Types (CCT) were identified using a Jensen-Shannon divergence matrix followed by a PCA then a BCA. Ellipses represent 90% confidence intervals. **E.** Genus and **F.** Bifidobacterium species relative abundance stratified by CCT. All statistical tests conducted are Mann-Whitney Wilcoxon tests corrected with FDR.

Using the Siamcat algorithm ^43^ we detected more than 40 taxa differing between infant and mothers at 2 years follow-up, from which *Coprococcus*, *Faecalibacterium*, *Dorea*, *Alistipes* and *Ruminococcus* were significantly more abundant in the mothers ; while *Bifidobacterium*, *Blautia*, *Anaerostipes* and *Veillonella* were significantly more abundant in the infants (Figure 3C). We validated the Siamcat results by running the ALDEx2 algorithm (aldex.ttest) which returned 24 significantly different genera (21 overlapping between the algorithms) (Supplemental figure 4A and Supplemental table 3). Focusing on the species level, we observed an increase of Bifidobacterium *breve*, *catenulatum* and *longum* while mothers had higher abundance of *Bifidobacterium adolescentis* (Supplemental figure 4A). Overall, the mother’s microbiota had higher relative abundance of Firmicutes and lower abundance of Actinobacteria (Supplemental figure 4B).

We then looked at the microbiota composition of the infants at 2 years solely and subjected the samples to an enterotyping approach^44^. Children at 2 years of age clustered into 3 enterotypes but not based on *Bacteroides*, *Prevotella* or *Ruminococcus* enterotypes as described in adults^44^ nor based on *Escherichia*, *Enterococcus* and *Staphylococcus* like for meconium microbiota. Child microbiomes, coined Child Community Types (CCT), clustered around *Ruminococcus*/*Blautia*/*Faecalibacterium (CCT1), Bifidobacterium (CCT2)* and *Clostridium sensu stricto 1*/*Escherichia*-*Shigella*/*Enterobacter (CCT3)* (Figure 3D). Additionally, we tested the relative abundance of the aforementioned genera and other discriminating genera (see above) between the Child community types showing that CCT 1 was discriminated by *Blautia*, *Faecalibacterium*, *Fusicatenibacter*, *Ruminococcus*, *Romboustia* and *Subdogranulum*; CCT 2 was discriminated mainly by *Bifidobacterium* and *Blautia*; CCT 3 was dominated essentially by *Bifidobacterium, Escherichia-Shigella, Clostridium sensu stricto 1* and *Enterobacter* (Figure 3E). Interestingly, CCT 2 and CCT 3 was dominated by different Bifidobacterium species as the first was dominated by *B. catenulatum* and the latter was dominated by *B. longum* (Figure 3F).

Then, we checked if the beta-diversity at 2 years was associated with any outcome at 1 year. Gut microbiota composition was associated with antibiotic usage during 1st year of life (PERMANOVA, p-value= 0.003, Supplemental figure 4C) but not with wheezing or infantile asthma or total allergies (data not shown). Antibiotic usage did impact the beta-diversity but not the alpha diversity (Supplemental figure 4C-D).

Maternal gut microbiota did not cluster in enterotypes driven by *Bacteroides*, *Prevotella* and *Ruminococcus* as described in Arumugam *et al* ^45^. Maternal gut microbiota clustered in Maternal community types (MaCT) around *Faecalibacterium*/*Prevotella*/*Coprococcus*, *Ruminococcus*/*Alistipes*/*Christensenellaceae R-7 group* and *Bifidobacterium*/*Blautia* for the MaCT 1 (54% of the mothers), MaCT 2 (29%) and MaCT 3 (17%) respectively (Supplemental figure 4E). While MaCT 1 and 2 resembled adult-like microbiomes, with a high relative abundance of Firmicutes and strict anaerobes, MaCT 3 was closer to infant microbiome composition (Supplemental figure 4F). Finally, alpha diversity analysis using Shannon entropy, showed strong differences in terms of diversity both between CCTs and MaCTs (Supplemental figure 4G).

### Breastfeeding has an impact on IgA bacteria binding at birth and was correlated to clinical outcome at 1 year

Purified stools were analyzed by flow cytometry using a 6 color panel targeting human IgA, IgM, IgE and IgG as well as bacterial Fc binding properties using a fluorescently labeled Fragment Crystallizable (Fc) region protein (Supplemental table 4). Bacteria cells were gated on size and structure to separate aggregates and debris from bacteria. Bacteria and the immune system interact *in vivo* and can produce aggregates of bacteria due to the dimeric sIgA. We therefore made a gating on the high structural events where we should find aggregates of bacteria. Each subsequent gating was made on unstained negative controls to account for auto-fluorescence in each channel (Figure 4A). A secondary validation was done using isotype controls to ensure that there was no residual staining (data not shown). Whereas anti-IgA, anti-IgM and anti-IgG antibodies were in F(ab’)2 format, the anti-IgE antibody was only available in whole-IgG antibody format, which required the removal of non-Fab mediated binding. We therefore evaluated IgE staining on Fc negative microbes to remove nonspecific staining due to Fc binding (Figure 4A). This analytical method could theoretically result in underestimation of IgE-binding, since certain microbes may simultaneously bind Fc and be targeted by IgE. Finally, we monitored the number of bacteria per gram of feces for each point in time. We found a median of 4.5×10^9^ bacteria per gram for the meconiums, which was significantly lower than the 1.2×10^10^ bacteria per gram found in feces from the 24 months old children as well as the mothers (Supplemental figure 5A). Importantly, this demonstrated that meconium contained enough microbes to confidently proceed with sequencing and IgAseq analysis.^46^

**Figure 4.**
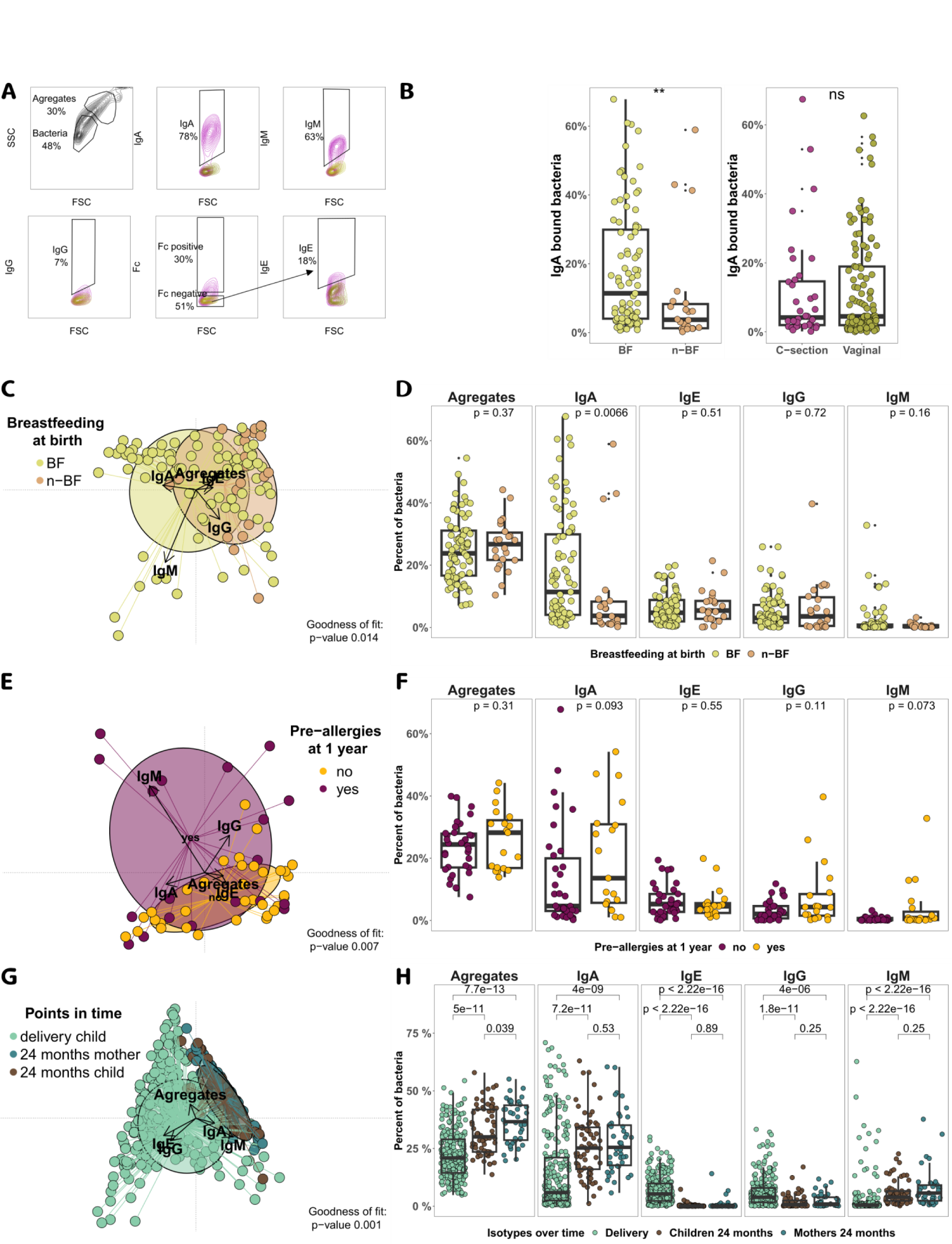
Antibody-microbiota phenotype at birth was linked to breastfeeding and pre-allergy outcomes later in life. **A.** Gating strategy realized on negative control (unstained control). Using Forward Scatter (FSC, represent size) and Side Scatter (SSC, represent granularity) we gated on bacteria and aggregates (large size and granularity). We then used the bacteria gate to quantify the percentage of IgA, IgG, IgM and Fc bound to bacteria. To quantify the percentage of bacteria bound to IgE we used the Fc negative gate to remove nonspecific staining. **B.** IgA bound bacteria according to breastfeeding (left panel) and birth route (right panel). Correspondence analysis of flow cytometry phenotype at birth on the right panel and boxplot of each gating on the right for breastfeeding (**C.** and **D.**), all allergies (**E.** and **F.**) and point in time (**G.** and **H.**). All statistical tests conducted are Mann-Whitney Wilcoxon tests.

We first analyzed the bacteria bound to IgA at birth. Meconium from breastfed neonates harbored significantly more IgA at the surface of bacteria than non-breastfed infants (median=11.4% and 3.7%, respectively, Mann-Whitney Wilcoxon test, p=0.006, Figure 4B). Birth route and MeCT however did not impact IgA levels at the surface of bacteria despite having strong differences in microbial composition (Figure 4B and Supplemental figure 5B). IgA was the isotype most represented at the surface of bacteria, followed by IgE, IgG and IgM (Supplemental figure 5C). Next we projected the five parameters using a correspondence analysis and fitted clinical factors to the regression. Breastfeeding resulted in a statistical difference mainly explained by IgA and IgM (Goodness of fit (GoF) = 0.002, Figure 4C). Breastfeeding did only impact the levels of IgA but not aggregates, IgE or IgG. IgM had a tendency to increase with breastfeeding but did not reach significance (Mann-Whitney Wilcoxon, p= 0.16, Figure 4D). Total allergies at 1 year, comprising infantile asthma, infantile wheezing and food allergy, was associated with the phenotypes at birth and driven by IgM and IgG (GoF, p= 0.033, Figure 4E). Infantile asthma and wheezing alone were also significatively associated with phenotypes at birth (GoF, p= 0.038 and p=0.07, Supplemental figure 5D-E). IgA levels were higher in future allergic infants but did not reach significance (Mann-Whitney Wilcoxon test, p=0.09, Figure 4F). Interestingly, infants with high levels of IgG and IgM at the surface of bacteria were all future allergic infants (Mann-Whitney Wilcoxon test, p=0.11 and p=0.073, Figure 4F).

We then compared the phenotypes between points in time and the mothers phenotypes. Interestingly, phenotypes are drastically different between meconium and later points in time (GoF, p=0.001, Figure 4G). These strong differences were explained by higher levels of aggregates and IgA but less IgE, IgM and IgG in the feces of children at 24 months and their mothers (Figure 4H). Additionally, staining levels are comparable between infants at 24 months and their mothers, except for aggregate levels, which were higher in the mothers (Mann-Whitney Wilcoxon test, p= 0.039, Figure 4H).

Maternal clinical factors, such as maternal allergies, were not linked to IgA levels at the bacterial surface in their neonates meconium (Supplemental figure 5F). Interestingly, IgE presence at the bacterial surface was neither associated with any perinatal factors (data not shown) nor mother clinical factors (Supplemental figure 5G). Finally, we tested immunoglobulins levels at the bacterial surface at 24 months in the children’s stools and found no association to any clinical factor (Supplemental figure 6).

### IgA binding to microbiota was still not fully adult-like at 2 years of age

To identify the gut microbes bound by IgA we performed magnetic cell sorting to separate IgA-bound microbes from IgA-non-bound microbes. The fractions were analyzed by 16S rRNA gene sequencing (IgASeq) and microbes with differential abundance between the two fractions were identified using paired statistical analysis (see IgASeq score section in M&M and Annexe 1), providing a novel IgASeq score for each identified taxa. This score is interpretable as a classic Z-score (IgA-bound microbes will have an IgAscore > 1.96 and IgA-non-bound microbes will have an IgAscore < −1.96 with a type-1 error of 5%). Gut microbiota with more than 10% IgA-bound bacteria were selected for IgAseq analysis. We detected 63 genera significantly enriched in the IgA^+^ fraction and 83 genera enriched in the IgA^−^ fraction (standalone ASV found only one time in either fraction were removed from further analysis). We have detected progressively more IgA-bound bacteria with increasing age (Figure 5A). Meconiums and 2 month samples were relatively sparse (median of 2.5 and 3 ASVs enriched in the positive fraction, respectively) and 24 months samples were more diversely enriched (Figure 5B). Although the IgA+ fraction was getting more diverse during the first 2 years of life, infants at 24 months had significantly less ASVs being targeted compared to mothers (Figure 5B). Additionally, the number of IgA^+^ and IgA^−^ enriched bacteria increased synchronously with overall gut microbiota alpha-diversity (Figure 5C, left and right panel, respectively). Meconium microbiota are dominated by *Escherichia*, *Enterococcus* or *Staphylococcus*. Interestingly all three taxa were bound to IgA at birth (Figure 5D), especially for *Staphylococcus* and *Enterococcus*. *Escherichia*-*Shigella* and *Bifidobacterium* were less targeted than the two other dominating taxa at birth (genera not different from zero, p-value > 0.05, Wilcoxon rank sum test) while at 2 months of age *Bifidobacterium* was less targeted and *Escherichia*-*Shigella* became extensively enriched in the IgA^+^ fraction (genera different from zero, p-value < 0.05, Wilocoxon rank sum test, Figure 5D). As new bacteria arose, the number of microbes bound to IgA equally increased. At 2 months *Veillonella* appeared and was enriched in the IgA^+^ fraction, while *Lactobacillus* and *Lactocaseibacillus* were not (Figure 5D). *Enterococcus faecalis* was frequently bound to IgA in gut microbiota at birth and at 24 months, while it was not enriched in the IgA^+^ fraction in maternal stool samples (Figure 5E). *Bacteroides* genus was consistently significantly enriched in the IgA^−^ fraction and detected at every point in time (Figure 5E). While *Bifidobacterium* and *Coprococcus* genera were generally enriched in the IgA^−^ fraction (Figure 5E and Supplemental figure 7), we saw differences at the species level. *B. bifidum* had a tendency to be enriched in the IgA^+^ fraction but failed to reach significance, while *C. comes* was strongly enriched in the IgA^+.^ fraction for the mother’s (Figure 5E). Finally, we saw differences between infants at 24 months and their mothers. Indeed, we could see more adult-like bacteria were enriched in the IgA^+^ fraction of the mothers than in the infants, such as *Coprococcus comes*, *Anaerostipes hadrus*, *Turicibacter* and *NK4A214 group* (Figure 5E and supplemental figure 7).

**Figure 5.**
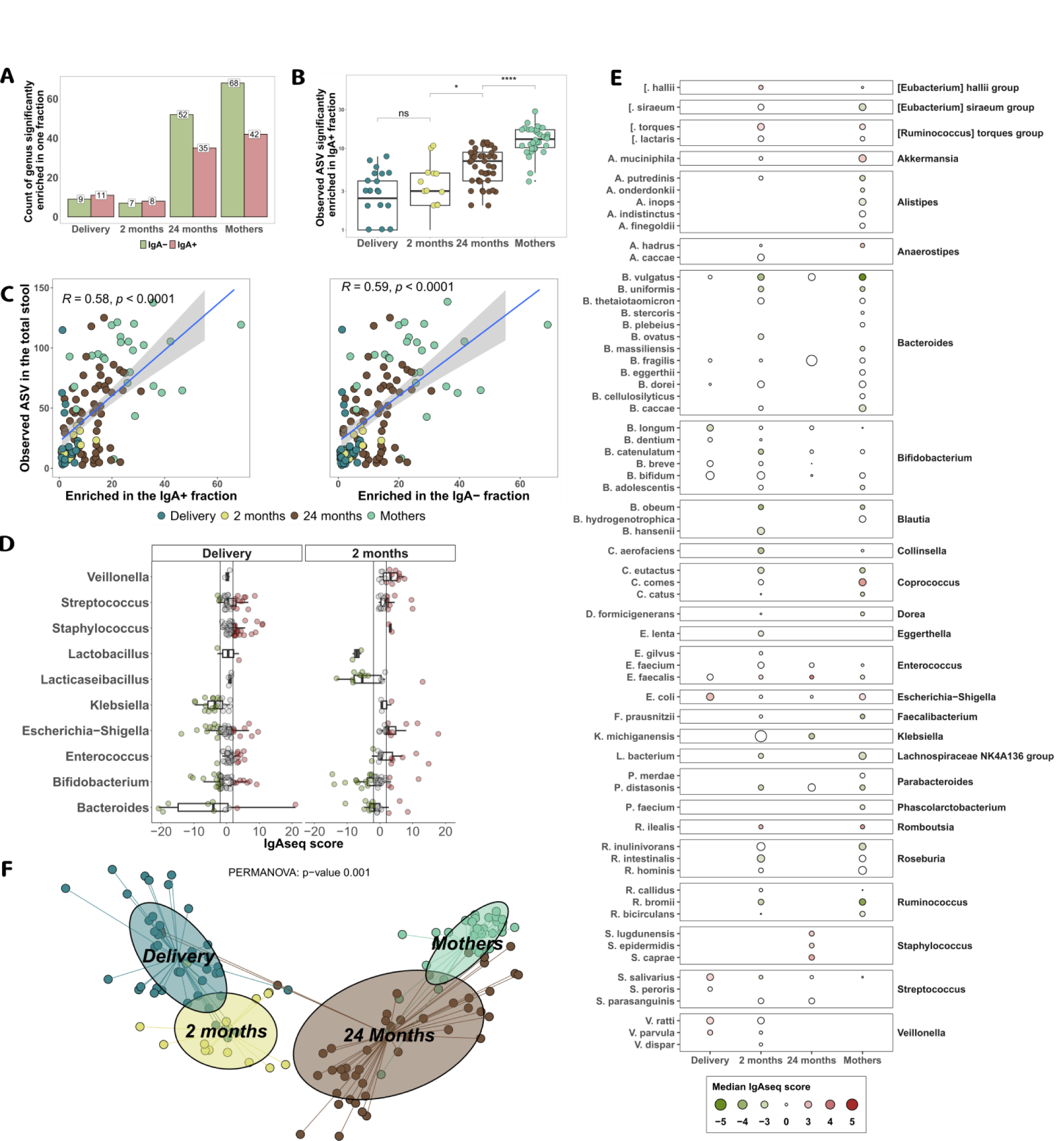
IgA binding to microbiota increases with new bacteria colonizing the microbiota. **A.** Total distinct genera detected as enriched in IgA-bound (IgAseq score > 1.96) or IgA-unbound (IgAseq score < −1.96) microbiota by points in time (at least detected in 2 samples). **B.** Number of distinct IgA-bound genera (ASVs) per sample by points in time. **C.** Correlation between observed ASVs in total stool and observed ASVs enriched in the IgA-bound (left panel) and IgA-unbound (right panel) microbiota. **D.** IgASeq score for delivery and 2 month samples at genus level. **E.** IgASeq scores for all points in time at the species level that were detected at least twice. Size of the dot represents the median absolute IgASeq score for each species. Green (IgAseq score < 0) and red (IgAseq score > 0) colors are associated with species significantly enriched in the IgA-unbound and IgA-bound microbiota, respectively. Color saturation reflects the p-value of the Wilcoxon rank sum test. **F.** PCoA of IgAseq score based on a Canberra distance for all points in time.

We were then interested in comparing the global IgAseq response trajectory over time using a PCoA to investigate dissimilarity and a PCA+BCA to find the most contributing taxa for individual time points. We subjected the IgAseq scores for individual taxa and donors to a PCoA based on a Canberra distance (adapted to values centered around 0) and segregated the samples by point in time. As presented in Figure 5F, IgAseq results for meconium and 2 month infant samples were very distant from children at 24 months and more so from mothers. 24-month-olds are different from their mothers according to the PCoA first and second axis (PERMANOVA, p-value =0.001, Figure 5F). In order to find the most discriminating taxa in the IgAseq dataset we selected the 225 ASVs enriched in the IgA^+^ fraction at least in one sample. We used this reduced IgAseq dataset to perform a PCA followed by a BCA grouped according to points in time. We then used the accumulated contribution to Axis 1 and 2 at the genus level (all ASV contributions grouped at the genus level, Supplemental figure 8A-B) and the ASV contribution to select the top 15 taxa contributing to each axis. Meconium samples were characterized by IgA+ enriched *Enterococcus*, *Staphylococcus* and *Streptococcus,* and overall lack of more mature taxa such as *Faecalibacterium*, *Akkermansia*, *Blautia* or *Ruminococcus* (Supplemental figure 8A and 8B). Child gut microbiota at 24 months were characterized mainly by IgA+ enriched *Faecalibacterium*, *Bifidobacterium*, *Lachnoclostridium*, *Erysipelaclostridium* and *Blautia* (Supplemental figure 8A-B). Finally, maternal gut microbiota were characterized by IgA+ enriched *Coprococcus* (*comes*), *Christensenellaceae R-7 group*, *Akkermansia* (*muciniphila)* and *NKA214 group* (Supplemental figure 8A-D).

### Microbiota at 24 months was not linked with pre-allergic outcome

Meconium microbiota was well associated with later pre-allergic outcomes, long before the symptoms arose (Figure 2E). We therefore tested the association between pre-allergic symptoms and the composition of 24 month-old child microbiota. Interestingly, microbial composition in the children’s stool at 24 months showed no association with pre-allergies (Figure 6A, PERMANOVA p=0.475), infantile asthma or infantile wheezing (data not shown). Additionally, the antibody-bound microbiota phenotype was not segregating children that developed pre-allergic symptoms by neither a multiparametric model (Figure 6B) nor single parametric models (Supplemental figure 6). IgASeq was the only approach which tended to segregate children with pre-allergic symptoms (Figure 6C, PERMANOVA, p=0.097). In conclusion, pre-allergies at 1 year were strongly associated with microbiota composition and antibody-bound microbiota phenotype at birth, but no association was observed at 24 months-of-age (Figure 6D).

**Figure 6.**
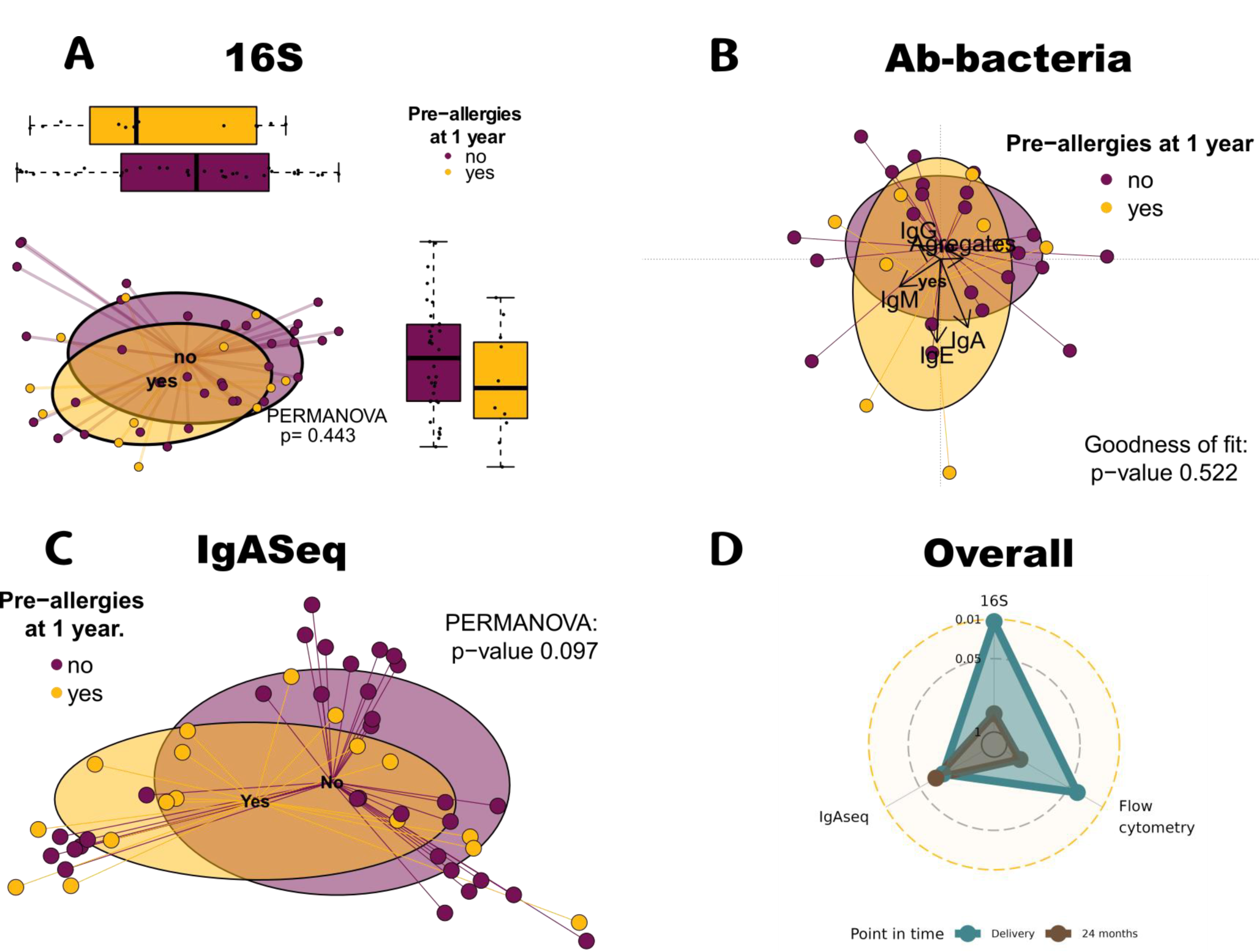
Microbiota at 24 months was not associated with allergy. **A.** PCoA of Bray-Curtis dissimilarity and **B.** Correspondence analysis of children antibody-bacteria phenotype at 24 months segregated by allergic status. **C.** IgASeq score of gut microbiota from 24 month-old children projected as PCoA and segregated by allergic status. A Canberra distance was used. **D.** Radar plot of the different technologies used to assess the association between allergies and the microbiota at different points in time. 16S refers to the total microbiota sequencing, Flow cytometry refers to antibody-microbiota phenotype projected with a correspondence analysis and IgASeq refers to the IgASeq score projected with a PCoA after a Canberra distance. P-values refers to the p-value presented earlier in the paper and in the present figure.

## Discussion

Meconium samples are very sparse in composition but are not sterile. We observed 10^10^ bacteria per gram of meconium covering a range of 10^8^ to 10^11^ bacteria per gram of meconium. Meconium microbiota therefore contains a microbiota biomass well within the analyzable range. Indeed, we have previously demonstrated that at least 10^6^ bacteria are required to obtain robust and representative 16S rRNA gene sequencing results ^46, 47^. Of note, this is 10-fold less than required for whole genome sequencing^48^. Meconium microbiota is very low in diversity and is mostly dominated by less than a handful of bacteria. By order of abundance we find *Escherichia*-*Shigella*, *Enterococcus*, *Staphylococcus*, *Bifidobacterium* and *Bacteroides.* Our results are in agreement with recent studies, showing a core meconium microbiome composed by *Escherichia*-*Shigella*, *Staphylococcus*, *Enterococcus, Streptococcus, Bifidobacterium* and *Bacteroides*^3, 8, 19, 25, 49–54^. Our findings and the mentioned recent literature are in contradiction with earlier work stating that meconiums were colonized by *Lactobacillus* and *Prevotella* for vaginally delivered children, while infants born by C-section were colonized by *Propionibacterium* and *Staphylococcus*^55, 56^. These discrepancies are probably due to the differences in sequencing technologies employed and the 16S region targeted. *Propionibacterium* and *Prevotella* are mainly detected in papers using 454 GLX Titanium sequencing^56, 57^, while the studies identifying the same consortia as us all used Illumina and 16S rRNA gene V3-V4/V4 region sequencing. Breastfeeding and birth route were the two main neonatal factors contributing to the microbial variations between neonates as previously described for the birth route.^54, 56–58^

We found no association between alpha diversity and clinical outcome at 1 year of age nor with maternal clinical status at delivery. On the contrary we found 3 meconium community types (MeCT), based on hierarchical clustering of meconium beta diversity and a direct association of microbiota composition at birth and pre-allergic symptoms. MeCT1 was dominated by *Escherichia*-*Shigella* genus, MeCT2 was diverse with no dominant taxa and MeCT3 was dominated by *Enterococcus*. These clusters were significantly associated with the birth route and almost reached significance for breastfeeding. MeCT1 was associated with breastfeeding and vaginal birth, while MeCT3 was more associated with formula feeding and C-section birth. Breastfeeding was associated with a lower usage of antibiotics during 1st year of life which is in line with previous studies showing breastmilk being protective against infections.^59^ Previous studies have identified associations between gut microbiota composition in 1 month old^60^ or 3-month-old infants^10^ and later clinical outcome, such as asthma or atopic wheeze. Here for the first time we demonstrate that meconium microbiota is linked to pre-allergic development. Additionally, we described three Meconium Community Types that are strongly associated with later clinical outcomes. MeCT 3 was associated with pre-allergic pathologies, infantile asthma and wheezing, while MeCT 2 was associated with food allergy and lactose intolerance.

It is well described that infant gut microbiota evolve rapidly in the first 2 years of life, with a particularly abrupt change associated with the transition from liquid to solid food^9, 24, 61^. In this context it is generally considered that the gut microbiota of a 2-3 years-old child has reached a fairly mature adult-like composition^62^. Here we demonstrate that the gut microbiota gradually evolves towards the maternal microbiota from 0 to 24 months of age. Interestingly, 2-year-old child gut microbiota remained compositionally less diverse than maternal gut microbiota, and we found numerous commensals associated with maternal gut microbiota, such as, *Ruminococcus*, *Coprococcus*, *Dorea*, *Faecalibacterium* and *Subdoligranulum*, being much less abundant in gut microbiota from 2-year-old children.

The fact that gut microbiota at birth and 3 months of life, but not at 12 or 24 months of life, are associated with pre-allergic clinical outcome later in life is somewhat contradictory. One possible explanation for this time gap between microbial exposure and clinical event could be the immunological nature of the relevant pre-allergic and probably later allergic diseases. Indeed, early-life immune maturation, priming and memory generation could have long-term clinical impact on the host. Indeed, clinical outcomes associated with the gut microbiota composition of 24-month-old children are all infectious in nature rather than immunological. Here we therefore propose to study the anti-microbiota immunological trajectories in children during the first 2 years of life. Most of the IgA found in the stool from breastfed infants during the first months of life are believed to come from maternal breast milk^63^. We show here that first pass stool (meconium) microbiota from breastfed infants are more frequently bound by IgA than non-breastfed infants. Of note, not every breastfed infant had IgA-bound gut microbiota and a few non-breastfed infants had a significant fraction of IgA-bound gut microbiota at birth. The latter result makes us propose that infants have a source of antibodies during perinatal life other than breastmilk. E.g. infants could gain immunoglobulins from amniotic fluid, which contain between 0.3 mg/mL and 0.7 mg/mL of IgA^64–68^ or IgA-bound microbes transferred horizontally from mother to child as well as from maternal bleeding during labor.

Interestingly, we also found free and microbe-bound IgE, as well as IgG and IgM, in both meconium and some 24 month-old child stools. While IgE has reportedly been found in feces of 1-year-old infants and adults^69, 70^, it is the first time IgE is found in the feces of meconiums both freely and bound to bacteria. However, we do not find any association with clinical factors at birth or later at 1 year both for the IgE in meconiums or at 2 years follow-up. We also considered that IgE levels in meconiums could be explained by maternal allergies, but we found no association with maternal clinical status at delivery. High levels of IgG and IgM in meconium were found only in neonates that developed pre-allergies later in life. Additionally, we describe the presence of IgG and IgM in the feces of children at 24 months and in their mothers. Even though the percentage of bacteria bound to IgG and IgM remains low at 2 years-of-age and in the mothers feces, we are (to our knowledge) the first to describe these immunoglobulin isotypes being present in the feces at steady state.

The interaction between microbes and antibodies are most often considered as opsonization. Opsonization is the process in which the Fab region of antibodies target antigens on pathogens, which can cross-link antigens and allows the free Fc region to promote the elimination of pathogens through various mechanisms, including complement activation and phagocytosis. However, some microbes, most often pathogens, express surface proteins that bind the Fragment crystallizable (Fc) region of antibodies, thereby efficiently reducing the host’s ability to eliminate the microbe^71^. Microbes expressing IgG Fc-binding proteins include *Streptococci* (protein G as well as M and M-like proteins^72–75)^, *Staphylococcus aureus* (protein A^76^). More recently it has been described that *R. gnavus* and *C. comes* express superantigens, which bind to the Fc region of immunoglobulins .^19^ Of note, the technology applied in our study does not allow us to differentiate between microbes bound to immunoglobulins through a normal antigen or a superantigen. Despite this limitation, multi-parametric flow cytometry phenotypic profiles of meconium microbiota were found to discriminate between breastfed and non-breastfed neonates, but also future pre-allergic children. However, phenotypic profiles of previously breastfed and non-breastfed 2-year-old children did not differ.

The number of taxa bound to IgA increases synchronously with alpha-diversity during the first 2 years of life. The IgA-bound taxa in children converges with the maternal IgA-bound taxa over time, but at 2 years of life major differences are still observed. Indeed, compared to mothers, children still had more frequent IgA-binding of *Faecalibacterium*, *Bifidobacterium* and *Lachnoclostridium* and less frequent IgA-binding to *Anaerostipes*, *Akkermansia* and *Ruminococcus*. We also found that intestinal IgA in breastfed children were targeting the main colonizing taxa *in vivo (Staphylococcus, Enterococcus* and *Streptococcus*). Our data are in accordance with Planer et al.^78^ Contrary to Dzidic et al., we more frequently found *Bacteroides* and *Roseburia* not bound to IgA, while *Staphylococcus*, *Streptococcus* and *Veillonella* were more frequently bound to IgA^79^. The specificity of gut IgA, as measured by IgASeq analysis, was the sole measure that tended to segregate pre-allergic and non-allergic children.

Earlier findings demonstrate an association between the gut microbiota at 1-3 months of life and later allergic outcome. Interestingly, this association was lost at 12 months of life^10, 42, 60^. Here we found an association between bacterial colonization immediately after birth (meconium) and the pre-allergic outcome later in life. It is important to note that little is known about pre-allergic signs in the neonates life. Infants with infantile asthma, food allergy and wheezing in the first year of life will not necessarily develop allergies later in life but have an increased risk of developing it^80^.

In our hands, birth route and breastfeeding were not directly associated with pre-allergic outcome, but they possibly impacted the meconium microbiota composition between an Escherichia, a mixed dominated or Enterococcus dominated meconium. The first is linked to a healthy outcome while the others are linked to an pre-allergic outcome. In concordance with previous studies, we did not see an association between microbial composition at 2 years of life and pre-allergic outcome. Our data therefore supports the notion that the first microbial encounter at birth may have long-term clinical impact on human health. We propose that this notion could be described as *“original microbial sin”*.

In conclusion, meconium composition is linked with pre-allergies later in life. We found that breastfeeding protects against infections. We also observed associations between antibody-microbiota phenotype and disease outcome both at birth and at 2 years of age. Maternal transfer of immunity via breastfeeding is essential in early life to ensure that gut microbiota is bound to IgA, which may regulate microbiota colonization in neonates. Although our results are purely associative, they support the notion that early-life interventions to change infant gut microbiota may have protective effects with regards to the onset of pre-allergic diseases later in life. Randomized interventional clinical trials should be envisaged to demonstrate the causality of early-life gut microbiota colonization and later onset of allergic diseases.

## Acknowledgments

We would like to acknowledge Catherine Lacluque, Diogo Griziotti, Martine Guinot, Laura Cerramon and David Codouel-Bovagnet who were implicated in the mother-child survey work. We finally acknowledge all the members of the EarlyFood consortium and the HEALS consortium (EU FP7-ENV Grant agreement ID: 603946). The project was funded by the JPI HDHL INTIMIC Era-Net call “Interrelation of the Intestinal Microbiome, Diet and Health”, Acronym: EarlyFOOD (https://www.healthydietforhealthylife.eu/76-hdhl-intimic-cofunded-call/396-earlyfood).

## Author contributions

Conceptualization and study design: R.V and M.L; HEALS/EXHES cohort conceptualization and study design: I.A.M., G.K.. Donor recruitment, sample collection and clinical data acquisition: I.A.M, G.K., M.S, C.L, J.R., L.C., G.P., S.L.G. and S.B.; Sample handling and storage: R.V., D.T., E.D., D.S., G.G., C.L. and M.L.; Data generation: R.V., D.T., E.D. P.F. and E.P; Data analysis, statistical analysis and data visualization: R.V. and M.L.; Manuscript writing: R.V. and M.L. All authors read, reviewed and approved the manuscript.

## Methods

### Cohort design

The HEALS (Health and Environment-wide Associations based on Large population Surveys) approach was applied to children from ten EU Member states drawn from the general population at maternity hospitals, who were recruited to an observational Exposure and Health Examination Survey (the EXHES Study)^81^. Biological bio-specimens sampled at birth were collected for a subset of mother-child dyads and as part of “The Long-term impact of 187 gestational and early-life dietary habits on infant gut immunity and disease risk” (EarlyFOOD) project^82^ we also followed some mother-child dyads longitudinally at 2 months and 24 months. 455 families from France were enrolled in the EXHES protocol in the Trousseau Hospital in Paris, France. Families were recruited during consultation for their pregnancy, 187 and 53 stool samples were collected for infants from birth and 24 months follow up, respectively (Supplemental figure 1). An additional 40 stool samples were retrieved from the mothers at 24 months. We also retrieved cord blood immediately after delivery and maternal blood samples the day after delivery. Additionally, we used 19 fecal samples from 2 months old infants derived from the Spanish EXHES cohort described in Rovira et al^83^. Mothers were requested to complete questionnaires at birth and 1 year after birth to procure mother-reported clinical information about mother and infant, including birth route and breastfeeding. For the mother we collected information about vaginal infection, gestational diabetes and allergy (asthma, rhinitis and food allergy). For the infant we collected information about sex, BMI, antibiotic treatment, otitis, bronchitis, diarrhea, cold, eczema and pre-allergic manifestations (wheezing, infantile asthma, food allergy). Pre-allergic symptoms were diagnosed by a medical professional for all infant asthma cases and 55% of wheezing cases.

### Stool processing

Raw meconium was collected at Trousseau hospital (Paris, France) and immediately cryo-preserved at −80°C . Follow up samples (24 months) were collected at home. Mothers were asked to collect a sample in a sterilized container and immediately freeze it at −20°C. Samples were transported to the laboratory where they were transferred to a −80°C freezer until processing. Raw stools were treated as previously described^46^. Raw stools were weighted and purified like the following : 200 mg of raw stools were aliquoted and reserved for DNA extraction and 16S rRNA gene sequencing while 1g of raw stool was reserved for immediate stool purification. Stool purification consisted of a series of centrifugation to retrieve semi-polar and polar components in fecal water and removal of debris. 1g of stool was weighted and homogenized in 5 mL of 1X Phosphate Buffer Saline (PBS Gibco) in a 50 mL tube. Tubes were then centrifuged at maximum speed (4600 rpm) for 20 min at 4°C. Supernatant containing polar and semi-polar components was retrieved and aliquoted for later use. Stool pellet was then homogenized in 50 mL of 1X PBS and centrifuged at 1000 rpm to pellet debris, 35 mL of supernatant are retrieved and centrifuged at maximum speed to pellet bacteria. Bacterial pellet is retrieved in 4 mL 1 X PBS 10% glycerol and frozen immediately at −80°C.

### Flow cytometry

Flow cytometry was performed on a Cytoflex S (Beckman Coulter) flow cytometer equipped with 4 lasers (405 nm, 488 nm, 561 nm and 638 nm) and 13 parameters. Purified stool was incubated with anti-human F(ab)^2^ IgA-FITC (Jackson research, 109-095-011), anti-human F(ab)^2^ IgM-DL405 (Jackson research, 109-476-129), anti-human F(ab)^2^ IgG-AF647 (Jackson research, 109-476-170), anti-human IgE PerCP-Cy5.5 (Biolegend, BLE325512) and human Fc biotin (009-060-008). After a wash step the latter was detected with PE-Cy7 conjugated streptavidin protein (AAT bioquest, 16917). The proportion of antibody-bound microbes was evaluated on a bacterial gate based on SSC-A/FSC-A to differentiate debris and microbial aggregates from bacteria. Background staining levels were evaluated with isotype control antibodies. Potential Fc-binding associated with the whole mouse IgG anti-human IgE antibody was corrected by analyzing IgE-binding only on the Fc-negative bacterial population. Bacteria count per gram of stool was assessed using the Cytoflex S flow cytometer. Briefly, Cytoflex S is capable of recording the amount of sample taken from the wells which is directly proportional to the number of bacterial cells present in the 1 microliter of purified stool used for the experiment. Reporting the number of bacterial cells calculated as a function of time and flow rate speed allows us to find the number of bacteria present in one milliliter of purified stool. Flow cytometry analysis was conducted with FlowJo software and R Bioconductor FlowCore package^84^.

### Immunoglobulin bound bacteria enrichment

100 μL (approx. 10^9^ microbes) of thawed purified stools were homogenized in 1 mL of 1X PBS and incubated with anti-human IgA-FITC antibody for 20 minutes at 4°C in the dark. Samples were washed in 1X PBS and incubated with 20 μL of anti-FITC magnetic beads (Miltenyi Biotech) for 20 minutes at 4°C in the dark. Samples were then enriched on magnetic columns using a multiMACS instrument (Miltenyi Biotech) according to the manufacturer guidelines. Enriched fractions were then centrifuged at maximum speed and frozen as dry pellets until further processing.

### DNA extraction and 16S rRNA gene sequencing

200 mg of raw stool or sorted samples were used to extract DNA using MagBead 96 Zymobiomic extraction kit according to manufacturer guidelines. Bead-beating step was performed on a Precellys 24 (Bertin) at maximum speed (7800rpm) for 60 seconds and repeated three times with one minute on ice, for a total of 3 minutes of cell lysing. DNA was retrieved in 50 μL of pure DNAse and RNAse free water and immediately stored at −20°C until amplification. The V3–V4 region of the 16S rRNA gene was amplified using a semi-nested PCR protocol. The semi-nested PCR was performed as previously described^46^. Briefly, 16S rRNA genes were amplified using a short pre-amplification step of 10 cycles with primer couple S-D-Bact-0341-b-S-17 (+ 341; 5′ CCTACGGGNGGCWGCAG 3′) and S-D-Bact-1061-a-A-17 (+ 1061; 5′ CRRCACGAGCTGACGAC 3′) (annealing temperature: 55 °C) followed by a second amplification with 30 cycles of standard PCR with the primer couple S-D-Bact-0341-b-S-17 (+ 341, 5’-AAGACTCGGCAGCATCTCCA-**CCTACGGGNGGCWGCAG**-3’) and S-D-Bact-0785-a-A-21 (+ 785; 5’-GCGATCGTCACTGTTCTCCA-**GACTACHVGGGTATCTAATCC**-3’) tagged according to the sequencing platform. Amplification reactions were performed with OneTaq HotStart (NEB, USA). After PCR purification, a dual index of 8 bases and P5-P7 adaptors were added by PCR with the primer couple 5’-AATGATACGGCGACCACCGAGATCTACAC-[i5]-ACACTCTTTCCCTACACGACGCTCTTCCGATCT-AAGACTCGGCAGCATCTCCA-3’ and 5’-CAAGCAGAAGACGGCATACGAGAT-[i7Rev]-GTGACTGGAGTTCAGACGTGTGCTCTTCCGATCT-GCGATCGTCACTGTTCTCCA-3’. After a new purification with Ampure beads, libraries were pooled in equimolar quantities. The final pool was sequenced on ILLUMINA Miseq (ICM, Paris, France) with a 2*300 V3 cartridge (2*25Millions of 300bases reads), corresponding to approximately a mean of 100 000 reads per sample after demultiplexing. The quality of the sequence run was checked internally using PhiX, and then each paired-end sequence was assigned to its sample with the help of the previously integrated barcodes.

### Bioinformatics for 16S rRNA gene analysis

Further processing of demultiplexed sequence reads followed the DADA2 in R software environment (version 4.2.3) and the DADA2 package (version 1.5.2)^85^. To ensure that amplicon sequence variants (ASVs) could be detected longitudinally between paired samples we performed a DADA2 pipeline with all the different runs together, including total stools for meconiums and follow up, along with IgAseq samples from both points in time. Runs (4 runs of MiSeq) were pooled together from the start and processed with the same parameters. Filtering parameters were set to 260bp and 250bp length and 2 and 3 for the expected errors, for the forward and reverse sequences respectively. Thirty-five bases were removed from the start of the sequences to ensure primer removal and subsequent taxonomic assignment. Dereplication step was performed with all samples from all runs to ensure ASV overlap between samples, then chimeras were removed using the DADA2 algorithm. Finally, ASVs were assigned to genus and species level using the SILVA database version 138.1. We validated that running all runs together did not introduce significant differences by comparing a subset of 20 samples that had been analyzed across at least 2 of the 4 sequence runs. The 20 samples were either analyzed all together or stratified by sequence run and the resulting abundance table was compared using alpha-diversity (Observed richness and Shannon entropy index), beta-diversity (Bray-Curtis dissimilarity) and ASV overlap. Raw sequence data will be made available at EBI upon peer-reviewed publication. Code and Rmarkdown for the DADA2 and validation will be made available upon peer-reviewed publication.

### IgAseq score

To identify IgA-bound microbes we make use of a paired analytical approach, which compares ASV abundances obtained from the analysis of microbiota fractionated into IgA-bound and IgA-unbound microbial communities (see experimental procedure above, Supplemental fig. 9.A). To identify within a microbiota, which ASVs are found significantly more abundant in the IgA-bound fraction compared to the IgA-unbound fraction (Supplemental fig. 9.B-C), we employed an analytical approach inspired by the ratio-intensity analysis frequently employed in paired OMICs analysis, such as microarrays^86^. Conceptually, this analytical method makes use of the fact that in OMICs type data it is expected that the vast majority of variables (e.g. gene transcription for transcriptomics) remain unchanged between the two conditions tested (output1 and output2). Therefore using the overall distribution of the log_2_ transformed ratio between output1 and output2 will be a good estimator of the underlying distribution of experimental fluctuation. The real differences between output1 and output2 would therefore represent the outliers of this distribution, which can be identified statistically. The experimental variation tends to be associated with the absolute intensity of the signal. Stratifying the outlier identification by the intensity of the signals represented by a log_10_ transformation of the product of output1 and output2 will therefore allow for better identification of outliers at the extremes of intensity (Z-transformation, Supplemental fig. 9.D-G).

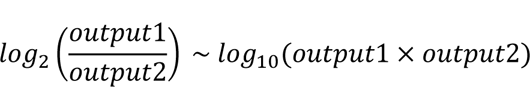

The underlying distribution of non-differential variables is well represented by this approach when the proportion of variables that are objectively different is low (e.g. a few hundreds out of approx. 20.000 genes), but if the proportion is higher and/or the number of variables smaller the conditions for this approach collapses. 16S rRNA gene analysis of gut microbiota will identify a few hundred ASVs in adult stool and much less in infants. Moreover, a significant part of these microbes will be bound by IgA. We have therefore altered the analytical strategy to accommodate these features. To establish the distribution of non-differential microbes we split the IgA-unbound fraction in two and treat both samples identically from DNA extraction to 16S rRNA gene sequencing (Supplemental fig. 9.C). The samples are biologically identical, which means that any variation between the samples can be considered due to experimental fluctuation. Using the experimental log_2_ transformed abundance ratio variation distribution as a function of log_10_ transformed abundance intensity (sliding-Z distributions) based on comparing the two IgA-unbound sequence results we can then assign an IgAseq score (Z score) to each ASV, which identify significantly differentially abundant ASVs for the comparison of IgA-bound and IgA-unbound microbiota (Z-transformation, Supplemental fig. 9.E and 9.G).

To facilitate this analysis we have created an R package “ImmuMicrobiome” (will be made available upon peer-reviewed publication).

### Data analysis

Bioinformatics and statistical analysis were performed in R (v4.2.3) using Rstudio IDE (2022.7.1.554). Microbiome composition was studied with the Phyloseq R package (v1.40). Statistical analysis and comparison of population distributions were performed using non-parametric Mann– Whitney Wilcoxon test, PERMANOVA, Goodness of fit and Chi Square tests. Plots were generated using ggstatsplot (v0.11.0), ggplot2 (v3.2.1), base R graphics and ade4 (v1.7.13). Figures were assembled using Inkscape software.

### Alpha-diversity analysis

Alpha-diversity analysis was performed using the Shannon diversity index using the Phyloseq R package (version 1.40.0). Additionally, we used the observed number of ASVs to assess the number of absolute ASVs detected in each sample.

### Differential abundance analysis

Differential abundance analysis was performed using two different approaches. We firstly used SIAMCAT (v2.0.1) function *check.association()* on genus relative abundance with a cutoff fixed at 0.001%. Since SIAMCAT is a recently published tool we associated these results with a conservative^87^ statistical test using ALDEx2 package (v1.28.1)^87^. Briefly, bacterial counts were normalized as centered log ratio (*aldex.clr()*), compared using the *aldex.ttest()* function and the *aldex.effect()* function. Results were then plotted using a volcano plot or bar plot visualization using ggplot2.

### Dimensionality reduction and ordination

Alpha-diversity was assessed as the Shannon diversity index and beta-diversity as the Bray-Curtis dissimilarity and subjected to a Principal Coordinate Analysis (PCoA) using the *dudi.pco* function from the ade4 package (1.7.22). Antibody-microbiota phenotype was projected using non constrained correspondence analysis with the *cca()* function provided by the vegan R package (2.6.4). Additionally, environmental factors were fitted to the data projection using the envfit() function in vegan. Principal Component Analysis (PCA) and Between Class Analysis (BCA) were performed as described in *Arumugam et al.*,^44^ except that we chose to use ASVs taxonomic level instead of genus level, for more details read the section “Clustering and enterotyping”. PCA and BCA were performed using the *dudi.pca()* and *bca()* functions package ade4 and plotted with ggplot2. PERMANOVA was used to assess statistical differences between projected groups using the Bray-Curtis dissimilarity and conducted in the vegan R package (2.6.4)^88^ with the *adonis2()* function.

### Clustering and enterotyping

In order to define clusters in the different microbiota at each point in time we used hierarchical clustering and enterotyping approaches. We assessed the performances of the enterotyping approach on meconium and projected the results using a PCoA according to the original paper^44^. However, meconium is very sparse and the enterotyping approach was less performing than a hierarchical clustering approach. Rapidly, the enterotyping approach produced overlapping clusters mainly due to the PAM clustering, while clustering with Ward.D2 produced well defined clusters when projected with a PCoA, these discrepancies were independent of the distance used. In that manner, we used the *hclust*() function and a Ward.D2 method to cluster meconium samples. For 24 month-old children samples and mother samples we used the enterotyping approach to compare our samples to previously described enterotypes. Both approaches gave between 1 to more than 10 possible clusters. To select an adequate number of clusters we used the Silhouette, WSS and Gap Stat algorithms thanks to the *fviz_nbclust()* function in the factoextra package (1.0.7)^89^. Finally, we validated the number of clusters with a PCoA projection using Bray-Curtis dissimilarity.

