## Supplemental information for "Gut microbiota and maternal immune transfer at birth influence pre-allergic clinical outcome"

EarlyFOOD study group: Catherine LACLUQUE[3], Laura CERRAMON[6], Giancarlo PESCE[5], Joaquim ROVIRA[4], Sandra BALDACCI[7], Bart KEIJSER[8,9], Jasper KIEBOOM[8], Stefania LA GRUTTA[10]

AFFILIATIONS:

1. Sorbonne Université, INSERM U1135, Centre d'Immunologie et des Maladies Infectieuses (CIMI-Paris), 75013, Paris, France

2. Département d'immunologie, Assistance Publique Hôpitaux de Paris (AP-HP), Hôpital Pitié-Salpêtrière, 75013 Paris, France.

3. Trousseau Maternity Hospital, Assistance Publique Hôpitaux de Paris (AP-HP), Sorbonne Université, Paris, France

4. Environmental Engineering Laboratory, Departament d'Enginyeria Quimica, Universitat Rovira i Virgili, Tarragona, Spain.

5. Institut Pierre Louis d'Epidémiologie et de Santé Publique, Equipe EPAR, Sorbonne Université, Paris, France.

6. Institute Desbrest of Epidemiology and Public Health, INSERM and Montpellier University, Department of Allergic and Respiratory Diseases, Montpellier University Hospital, Montpellier, France.

7. Institute of Clinical Physiology (IFC), National Research Council (CNR), Pisa, Italy

8. Department of Microbiology and Systems Biology, TNO Healthy Living and Work, Leiden, The Netherlands.

9. Department of Preventive Dentistry, Academic Center for Dentistry Amsterdam, University of Amsterdam and Vrije Universiteit Amsterdam, Amsterdam, The Netherlands.

10. Institute of Translational Pharmacology (IFT), National Research Council (CNR), Palermo, Italy.

Running TItle: Original microbial sin - First microbial colonizers are associated with pre-allergic manifestations in infants.

*Correspondence and requests for reprints should be addressed to:

Martin Larsen, Ph.D.,

Sorbonne University, Inserm UMR-S1135

Centre d'Immunologie et des Maladies Infectieuses (CIMI Paris), 5eme etage (bureau 508)

91 boulevard de l'Hôpital

75013 Paris

France

### Supplemental information

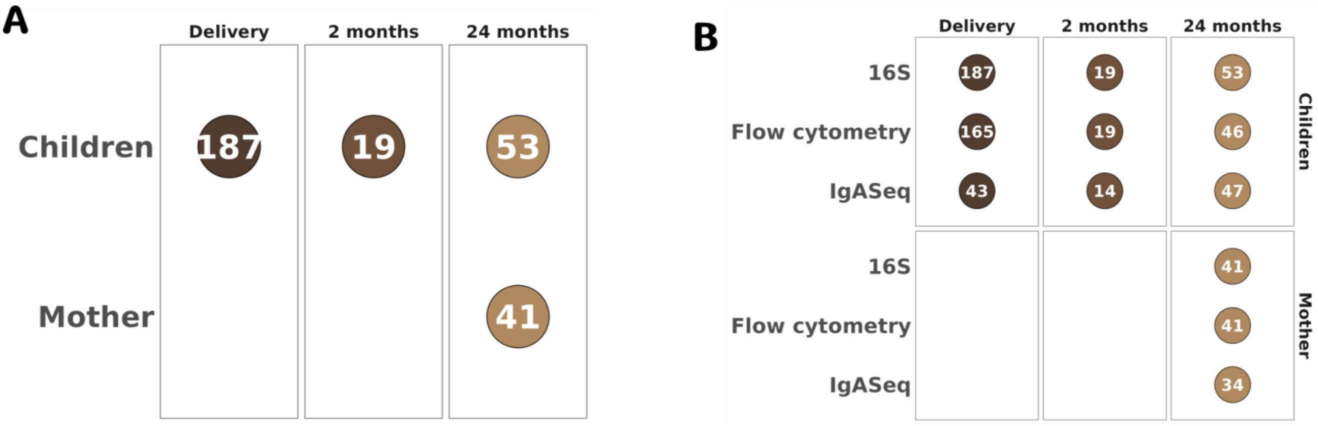


**Supplemental figure 1. EXHES cohort description.**

**A.** Description of the number of mother and child samples used at different points in time. **B.** Description of number of mother and child samples used for each technology.

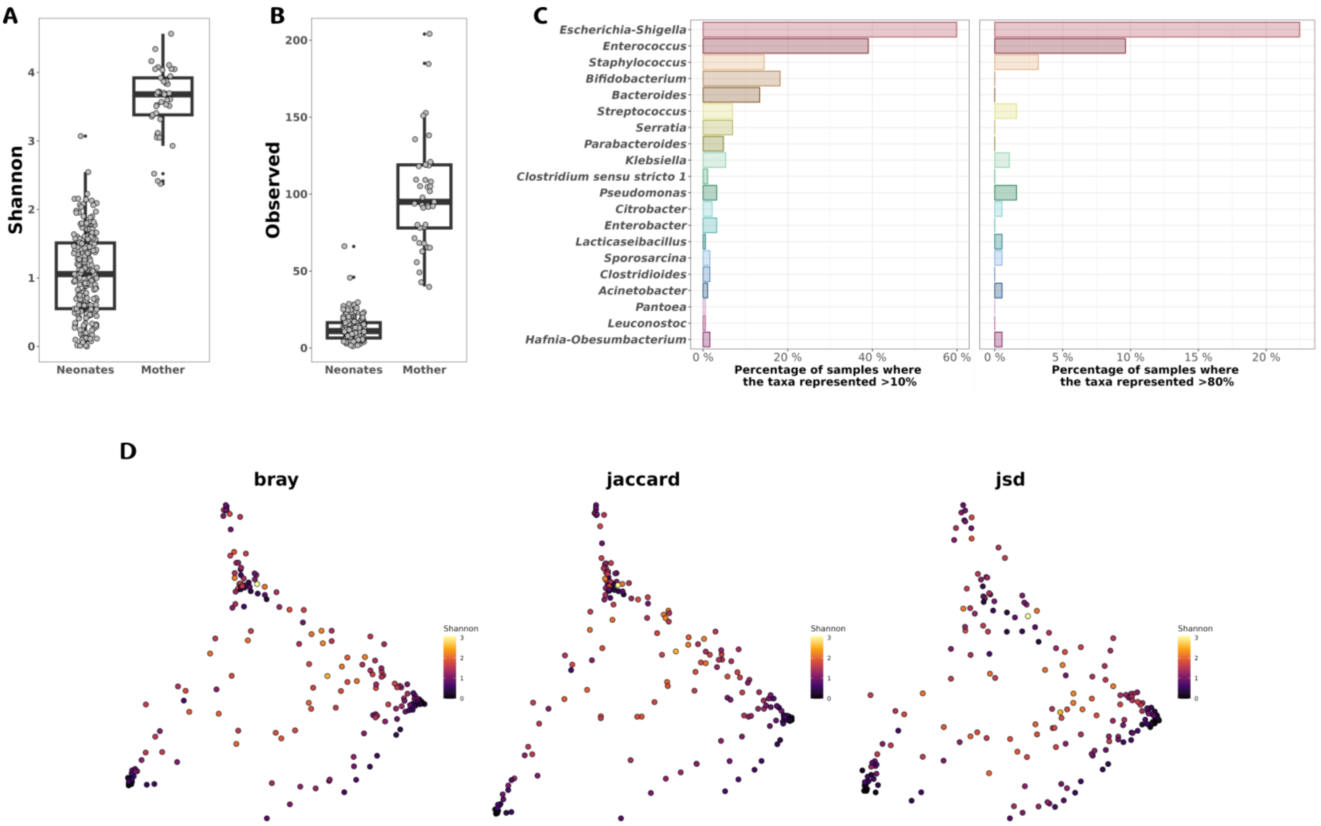


**Supplemental figure 2. Meconium microbiota was sparse and strongly dominated by a handful of taxa.**

**A.** Shannon alpha diversity for the meconium microbiota compared to their mothers. **B.** Observed number of ASVs for the meconium microbiome compared to their mothers. **C.** Percentage of samples where dominant taxa represent more than 10% (left panel) and 80% (right panel). **D.** Three distance matrices were computed with a PCoA and colored by Shannon index, from left to right, Bray-Curtis dissimilarity, Jaccard dissimilarity and Jensen-Shannon divergence.

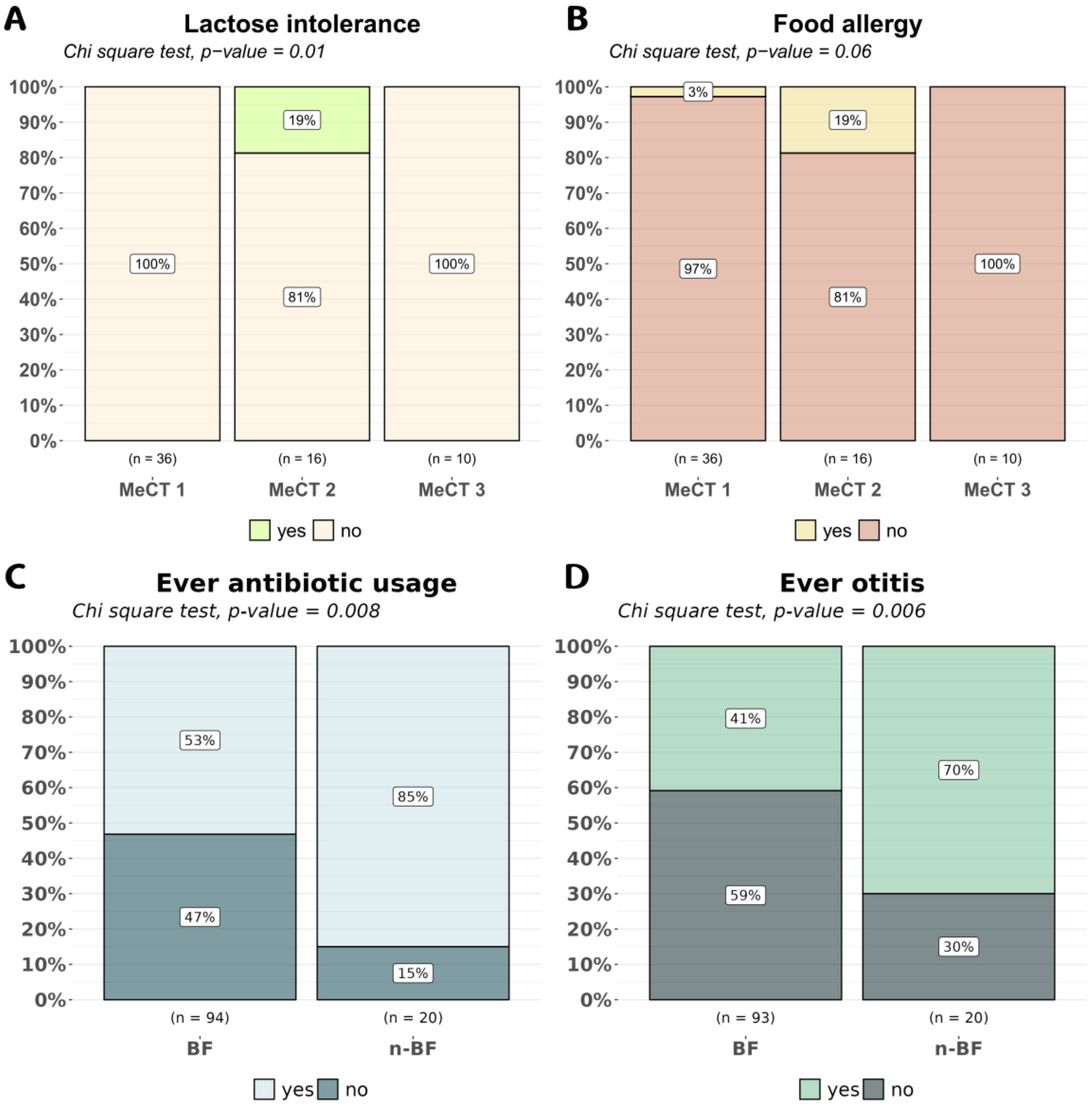


**Supplemental figure 3. MeCT are linked to allergies while breastfeeding protects against antibiotic usage and otitis during 1st year of life.**

Association between MeCT and **A.** lactose intolerance and **B.** food allergy. **C.** Association between breastfeeding and antibiotic usage during 1st year of life. **D.** Association between breastfeeding and otitis during 1st year of life.

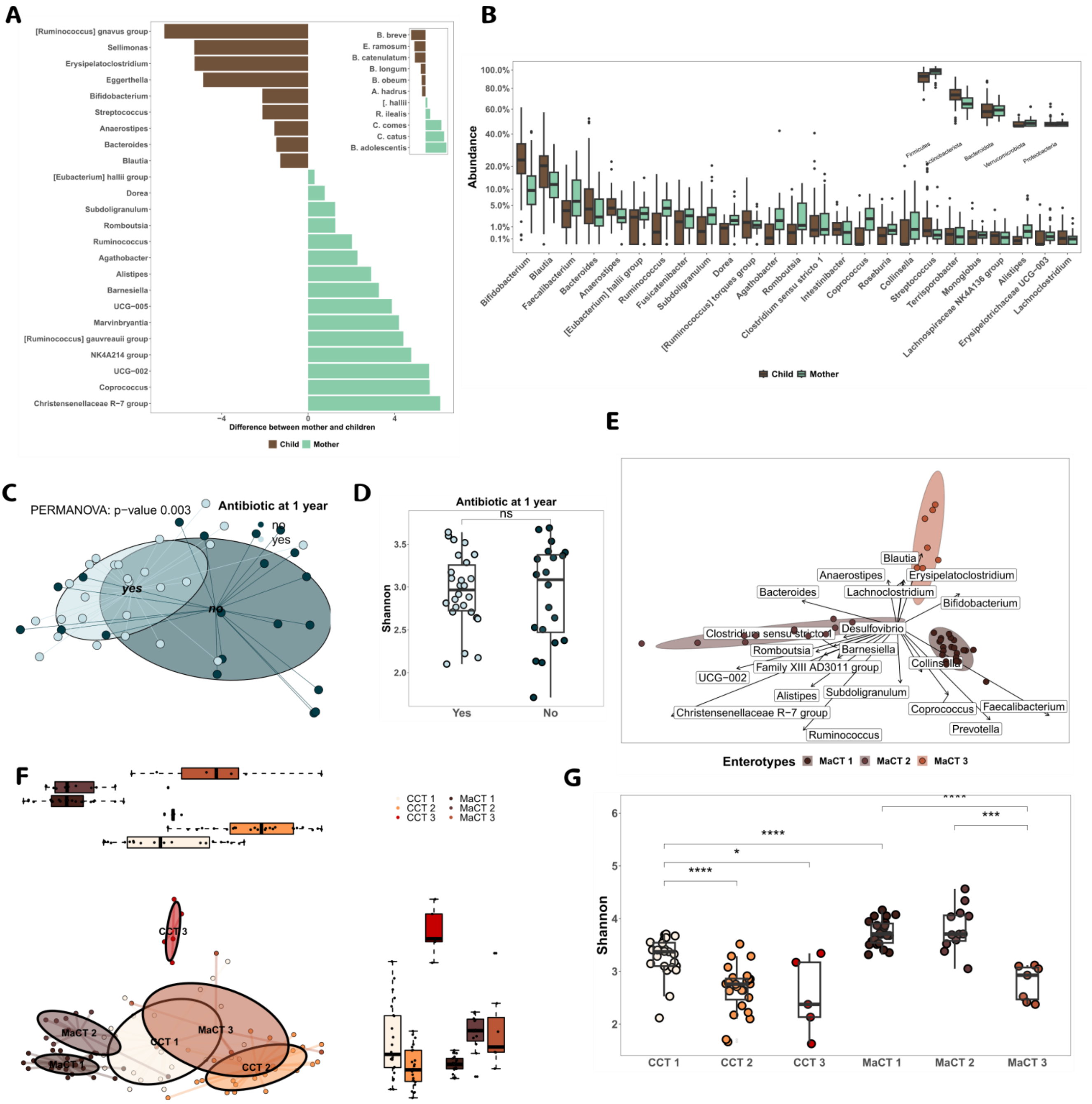


**Supplemental figure 4. Gut microbiota from mothers and children remain different 2 years after birth.**

**A**. ALDEx2 differential abundance analysis between mothers and children (2 years of age) gut microbiota at the genus level. All taxa depicted are significantly higher in either mothers (green) or children (brown). **B**. Boxplot of relative abundance for each genus, sorted by cumulated abundance and colored by groups. **C**. PCoA of child microbiomes alone and segregated by antibiotic usage during the 1st year of life. **D**. Impact of antibiotic consumption on gut microbiota alpha diversity at 2 years of life. **E.** Enterotyping of mother microbiomes merge at the genus level. A Bray-Curtis dissimilarity was used, followed by a PCA and a BCA. Taxa displayed are the top 15 genera contributing to each axis 1 or 2. **F.** Comparison of MaCT and CCT using a PCoA after a Bray-Curtis dissimilarity analysis. **G.** Comparison of alpha diversity of CCT and MaCT clusters. All statistical tests conducted are Mann-Whitney Wilcoxon tests corrected with FDR.

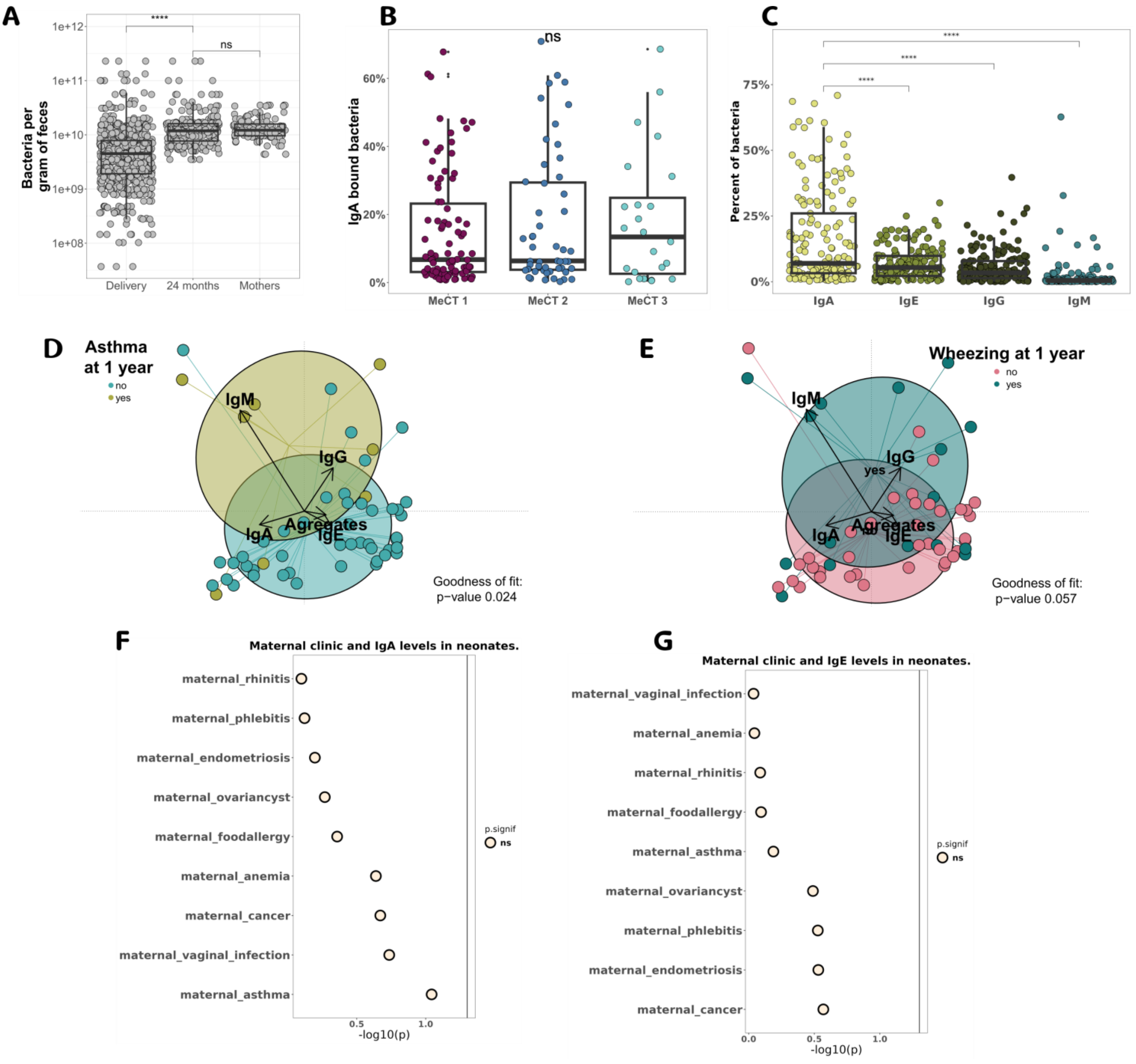


**Supplemental figure 5. Antibody-microbiota phenotype at birth was not linked with maternal clinical factors but to later pre-allergy outcome.**

**A.** Flow cytometry bacterial count for all points in time. **B.** IgA bound bacteria according to MeCT defined using 16S composition of the meconium samples. **C.** Overall distribution of antibody isotypes across meconium microbiota samples. Correspondence analysis of phenotypes at birth and segregated by **D.** asthma and **E.** wheezing at 1 year. Association between **G.** IgA- and **H.** IgE-bound bacteria and maternal clinical factors. All statistical tests conducted are Mann-Whitney Wilcoxon tests.

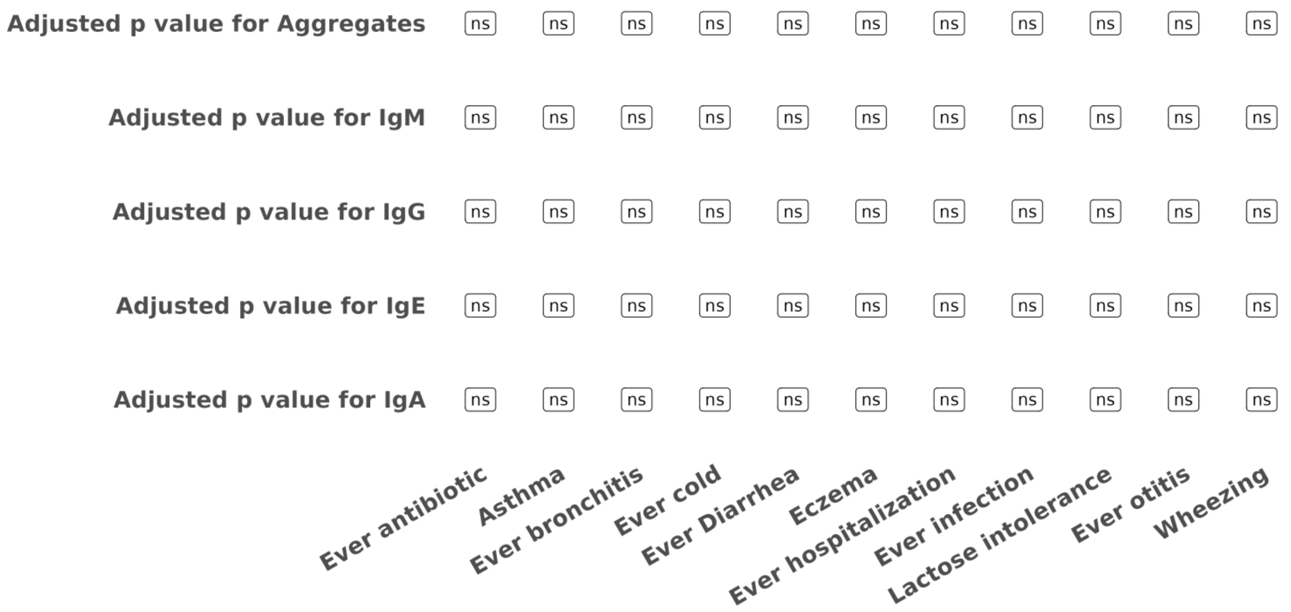


**Supplemental figure 6. Association between flow cytometric antibody-microbiota phenotypes for child gut microbiota at 24 months and clinical factors at 1 year.**

Antibody-microbiota phenotype analyzed for gut microbiota from children at 24 months of age (IgA, IgG, IgM, IgE and Fc-binding). Associations with clinical factors were conducted with an FDR correction Wilcoxon test.

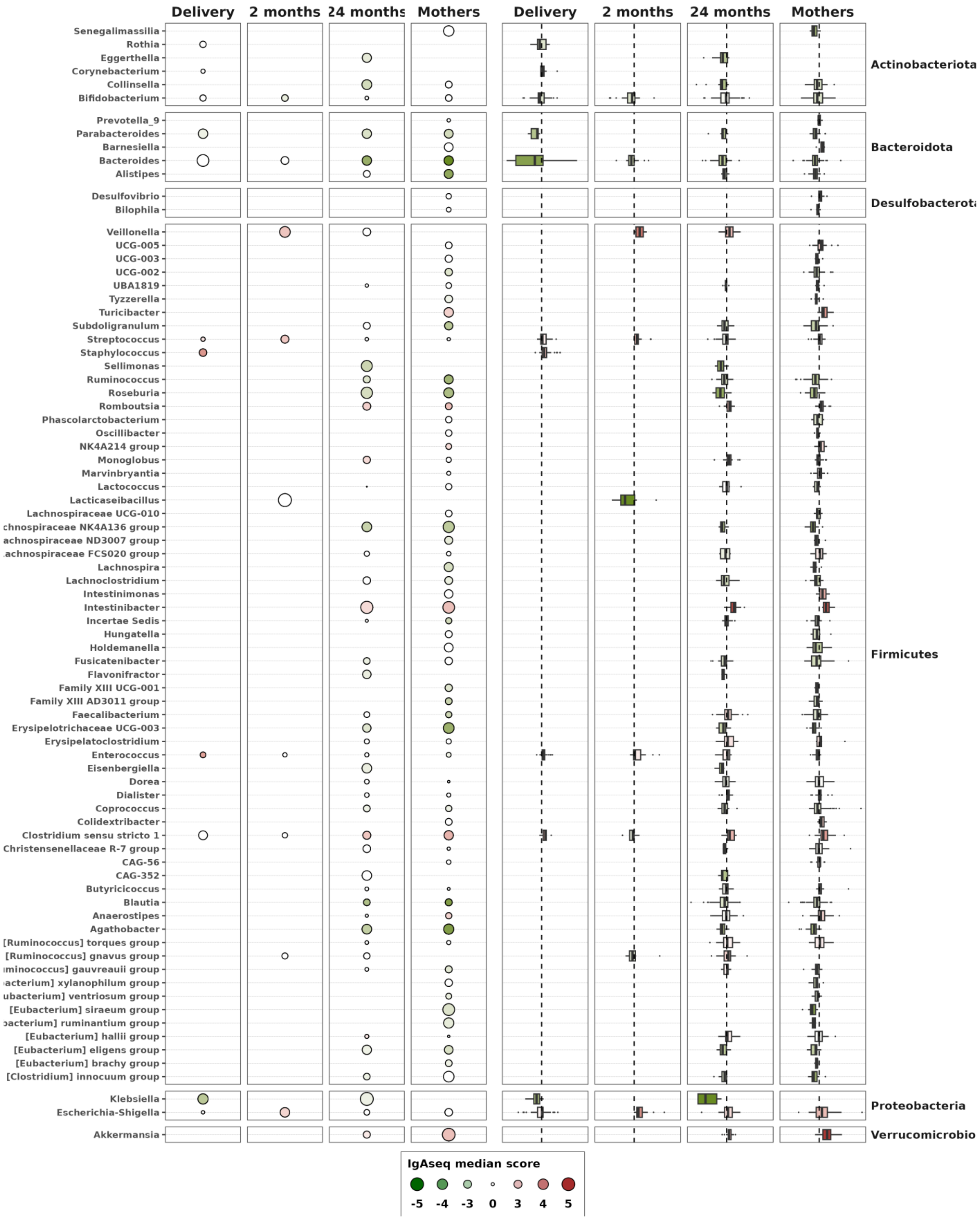


**Supplemental figure 7. IgASeq response at the genus level.**

IgASeq scores for all points in time at the genus level that were composed of at least two ASVs. **Left panel:** Size of the dot represents the median absolute IgASeq score for each species. Green (IgAseq score < 0) and red (IgAseq score > 0) colors are associated with genera significantly enriched in the

IgA-unbound and IgA-bound microbiota, respectively. Color saturation reflects the p-value of the Wilcoxon rank sum test. **Right panel:** Distribution of each IgAseq score data point is represented as box plots.

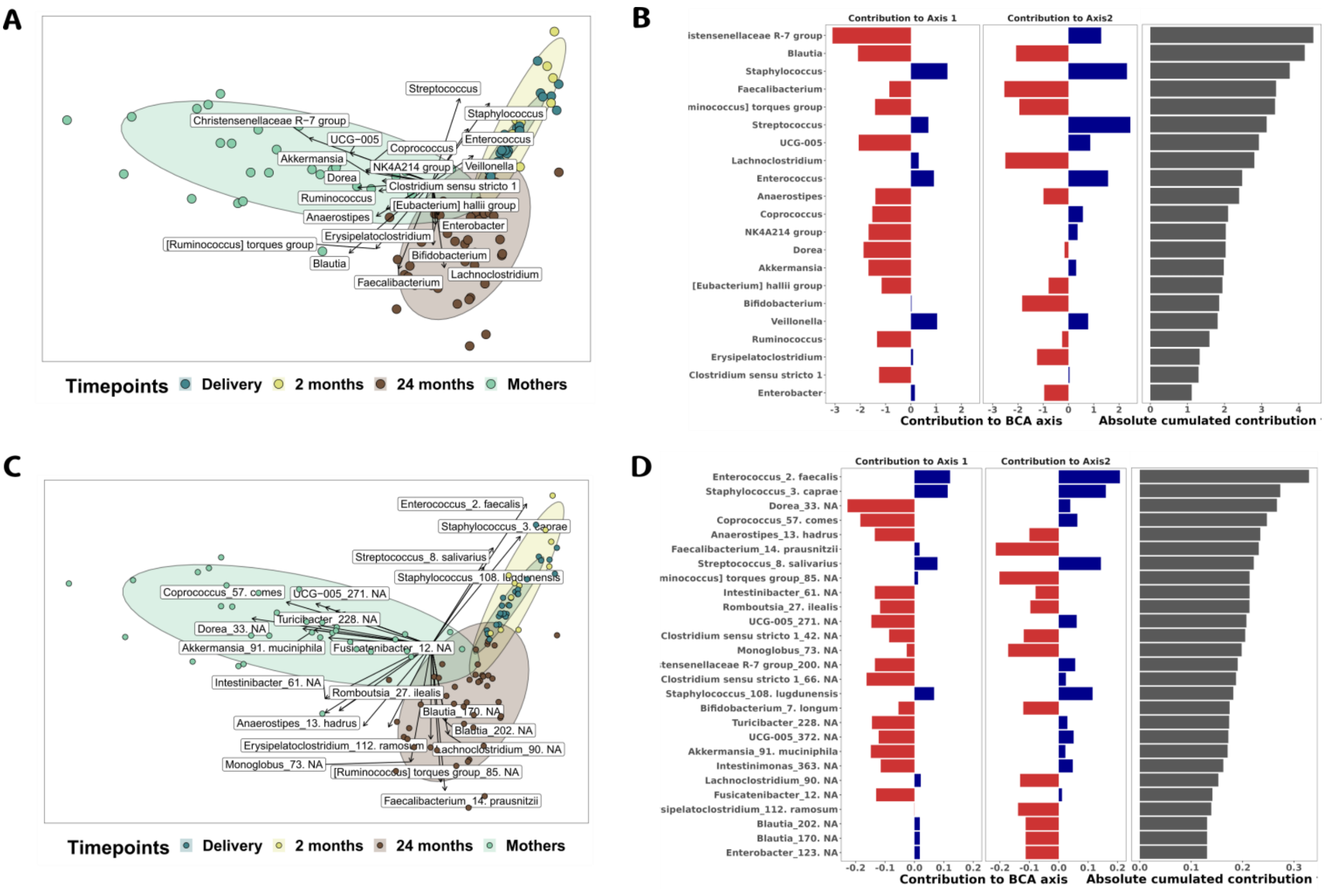


**Supplemental figure 8. IgASeq projection with PCA and BCA shows bacterial contribution to each point in time.**

PCA followed by a BCA on IgAseq-score. ASVs were trimmed to obtain all ASVs that were positively enriched in the IgA-bound microbiota at least once. **A.** Taxa contribution to axis were summed at the genus level **B.** Taxa contribution to axis depicted as bar plots and ordered according to their absolute cumulated contribution to BCA at genus level **C.** Taxa contribution to axis were summed at the ASV level. **D.** Taxa contribution to axis depicted as bar plots and ordered according to their absolute cumulated contribution to BCA at ASV level.

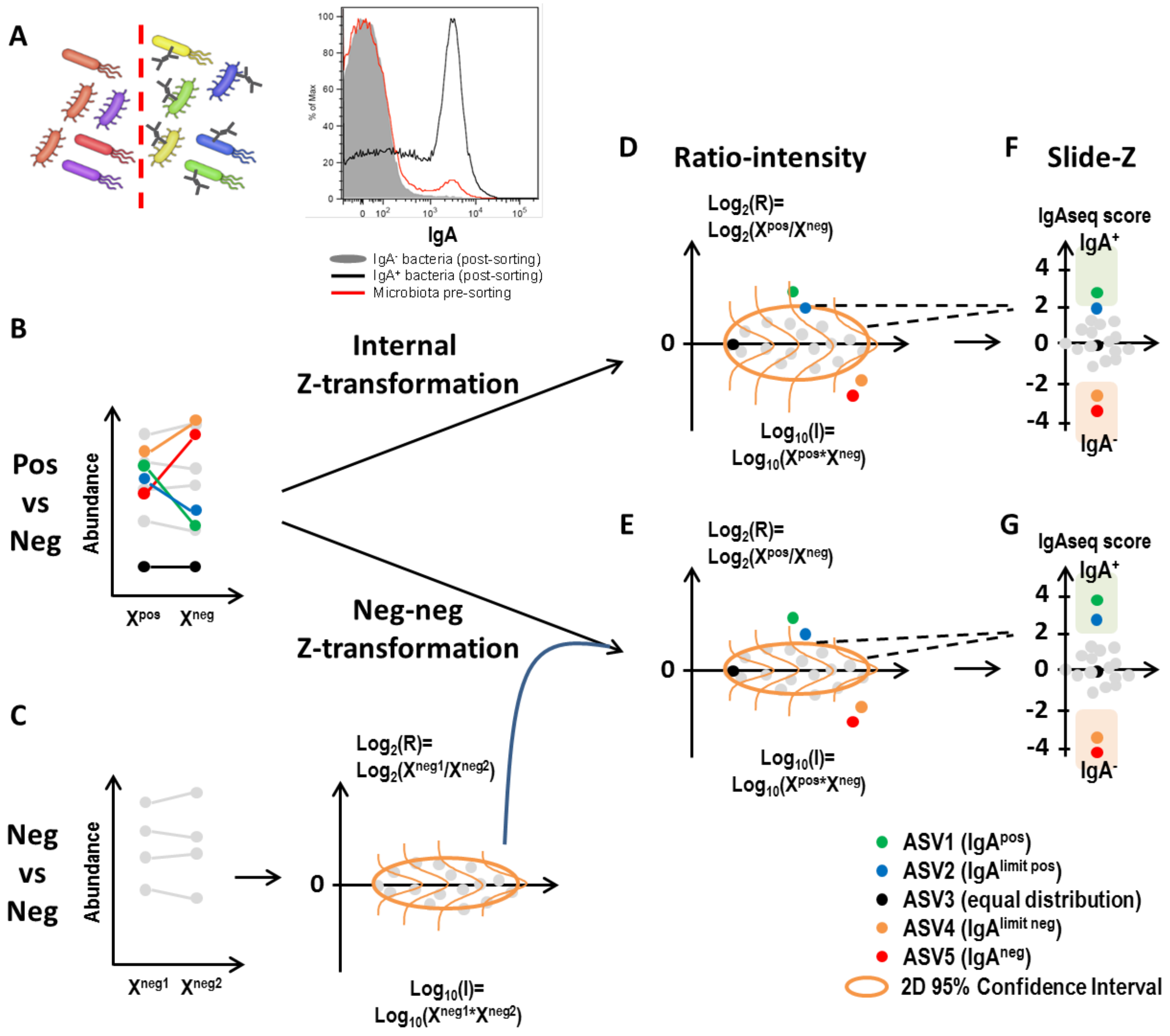


**Supplemental figure 9. Principles of IgAseq analysis based on Normal vs Neg-Neg Z transformation theory.**

**A.** Schematic representation and flow cytometric analysis of IgA-bound microbiota pre and post sorting. **B.** ASV abundance in IgA-bound ($X^{pos}$) vs IgA-unbound ($X^{neg}$) microbiota (biological variation) and **C.** ASV abundance in IgA-unbound ($X^{neg1}$) vs IgA-unbound ($X^{neg2}$) microbiota (experimental variation). **D.** $Log_2$ transformed abundance ratio plotted as a function of $log_{10}$ transformed abundance intensity for an IgA-bound (green), limit IgA-bound (blue), limit IgA-unbound (orange) and IgA-unbound (red) ASV overlapped with a 2-Dimensional 95% confidence interval. **E.** The same plot as D but overlapped with a 2-Dimensional 95% confidence interval derived from the distribution of repeated analysis of IgA-unbound microbiota (see C). **F.** and **G.** IgAseq score calculated as the sliding Z-transformation of the log2 transformed abundance ratio.

### Supplemental tables.

| Clinical factor | No | Yes | Adjusted p-value | Time of recording |
|---|---|---|---|---|
| Ever Antibiotic usage | 26 | 44 | 0.92 | 12 months |
| Asthma | 61 | 9 | 0.74 | 12 months |
| Ever Cold-Nasopharyngitis | 1 | 69 | 0.76 | 12 months |
| Ever Diarrhea | 19 | 51 | 0.92 | 12 months |
| Eczema | 50 | 20 | 0.77 | 12 months |
| Food allergy | 65 | 5 | 0.74 | 12 months |
| Hospitalization | 52 | 18 | 0.77 | 12 months |
| Lactose intolerance | 67 | 3 | 0.92 | 12 months |
| Maternal anemia | 138 | 25 | 0.77 | Delivery |
| Maternal asthma | 148 | 18 | 0.74 | Delivery |
| Maternal cancer | 162 | 1 | 0.92 | Delivery |
| Maternal endometriosis | 156 | 10 | 0.74 | Delivery |
| Maternal food allergy | 160 | 6 | 0.8078 | Delivery |
| Maternal ovariancyst | 144 | 21 | 0.92 | Delivery |
| Maternal phlebitis | 165 | 2 | 0.74 | Delivery |
| Maternal rhinitis | 149 | 17 | 0.77 | Delivery |
| Maternal vaginal infection | 125 | 40 | 0.74 | Delivery |
| Ever Otitis | 33 | 36 | 0.77 | 12 months |
| Pediatrician consultation | 10 | 60 | 0.74 | 12 months |
| PMI consultation | 48 | 21 | 0.77 | 12 months |
| Urgence consultation | 40 | 30 | 0.77 | 12 months |
| Wheezing | 49 | 21 | 0.74 | 12 months |

**Supplemental table 1.** Meconium Shannon alpha diversity tested for clinical factors at delivery and at 1 year. Wilcoxon test corrected with FDR.

| Factor | Pre-allergic (n=24) | Non pre-allergic (n=38) | p-value | p.signif | Clinical source |
|---|---|---|---|---|---|
| Birth route (% vaginal) | 75.0% | 68.4% | 0.704 | ns | Child |
| Breastfeeding | 75.0% | 73.7% | 1.000 | ns | Child |
| Eczema | 12.5% | 13.2% | 1.000 | ns | Child |
| Ever antibiotic | 79.2% | 52.6% | 0.066 | ns | Child |
| Ever bronchitis | 75.0% | 23.7% | 2.10E-04 | *** | Child |
| Ever cold | 100.0% | 97.4% | 1.000 | ns | Child |
| Ever hospitalized | 41.7% | 15.8% | 0.049 | * | Child |
| Ever otitis | 62.5% | 44.7% | 0.316 | ns | Child |
| Sex (% Female) | 37.5% | 47.4% | 0.491 | ns | Child |
| Maternal asthma | 8.3% | 13.2% | 0.820 | ns | Mother |
| Maternal food allergy | 0.0% | 2.6% | 1.000 | ns | Mother |
| Maternal gestational diabetes | 4.2% | 5.3% | 1.000 | ns | Mother |
| Maternal gluten intolerance | 4.2% | 7.9% | 1.000 | ns | Mother |
| Maternal rhinitis | 8.3% | 10.5% | 1.000 | ns | Mother |
| Maternal vaginale infection | 16.7% | 21.1% | 0.864 | ns | Mother |

**Supplemental table 2.** Distribution of infant and maternal clinical factors in infants with (n=24) and without (n=38) known pre-allergic manifestations at 1 year of age. The homogeneity of clinical factor distribution in allergic and non-allergic infants was tested with Chi-square tests.

| Genus | p adjusted SIAMCAT | Fold change | p adjusted ALDEx2 | Difference between ALDEx2 |
|---|---|---|---|---|
| Agathobacter | 1.8E-04 | -2.0E+00 | 9.5E-12 | -3.3E+00 |
| Alistipes | 4.8E-05 | -1.7E+00 | 1.6E-09 | -4.3E+00 |
| Anaerostipes | 9.6E-03 | 2.3E-01 | 4.1E-01 | 9.9E-01 |
| Barnesiella | 8.8E-05 | -1.8E+00 | 1.0E-02 | -1.3E+00 |
| Bifidobacterium | 4.9E-04 | 3.9E-01 | 6.4E-03 | 1.3E+00 |
| Blautia | 4.0E-02 | 1.6E-01 | 7.5E-02 | 3.9E-01 |
| Colidextribacter | 5.2E-04 | -1.1E+00 | 4.4E-03 | -1.7E+00 |
| Coprococcus | 5.6E-06 | -2.6E+00 | 4.9E-30 | -8.5E+00 |
| Dialister | 7.0E-03 | -1.3E+00 | 2.5E-02 | -1.6E+00 |
| Dorea | 1.8E-05 | -1.3E+00 | 5.8E-23 | -3.1E+00 |
| Eggerthella | 3.5E-03 | 1.5E+00 | 1.3E-07 | 3.0E+00 |
| Erysipelatoclostridium | 1.4E-04 | 1.8E+00 | 7.9E-11 | 4.2E+00 |
| Faecalibacterium | 1.2E-02 | -5.0E-01 | 8.7E-06 | -9.8E-01 |
| Intestinimonas | 1.1E-02 | -7.5E-01 | 8.1E-03 | -1.4E+00 |
| Marvinbryantia | 2.4E-06 | -1.8E+00 | 8.3E-13 | -5.3E+00 |
| Phascolarctobacterium | 3.7E-04 | -1.3E+00 | 1.1E-06 | -3.4E+00 |
| Romboutsia | 6.4E-04 | -1.3E+00 | 9.7E-21 | -3.9E+00 |
| Ruminococcus | 3.5E-04 | -1.9E+00 | 3.5E-21 | -4.5E+00 |
| Sellimonas | 6.4E-04 | 1.7E+00 | 5.9E-12 | 4.7E+00 |
| Subdoligranulum | 1.9E-03 | -1.7E+00 | 3.8E-22 | -7.4E+00 |
| UCG-002 | 1.1E-06 | -2.5E+00 | 5.6E-19 | -6.4E+00 |
| UCG-003 | 3.1E-03 | -5.1E-01 | 3.0E-02 | -1.1E+00 |
| UCG-005 | 3.9E-05 | -1.9E+00 | 1.2E-17 | -5.8E+00 |
| Veillonella | 1.2E-02 | 9.4E-01 | 2.2E-08 | 2.6E+00 |

**Supplemental table 3.** Overlapping taxa between SIAMCAT and ALDEx2 differential abundance analysis. Positive score indicates a higher abundance in the children (24 months old), negative score indicates a higher abundance in mothers.

| Target | Conjugate | Molecule structure | Mono- or polyclonal | Manufacturer | Cat. Number |
|---|---|---|---|---|---|
| IgA | FITC | F(ab)2 | Polyclonal | Jackson Immuno | 109-096-011 |
| IgG | Alexa Fluor 647 | F(ab)2 | Polyclonal | Jackson Immuno | 109-605-098 |
| IgM | DyLight 405 | F(ab)2 | Polyclonal | Jackson Immuno | 109-476-129 |
| IgE | PerCP-Cy5.5 | Whole IgG | Monoclonal | BioLegend | 325512 |
| Fc-binding domains | Biotin | Fc | Polyclonal | Jackson Immuno | 009-060-008 |
| Biotin | PE-Cy7 | Streptavidin | NA | AAT Bioquest | 16917 |
| DNA | NA | Syto X Orange | NA | Invitrogen | S11368 |

**Supplemental table 4.** Antibody list used for microbiota phenotype flow cytometry panel
